# Dynamical model and geometric insights in the discontinuity theory of immunity

**DOI:** 10.1101/2025.07.16.663927

**Authors:** Christian Mauffette Denis, Victoria Mochulska, Maya Dagher, Vincent Verbavatz, François X.P. Bourassa, Grégoire Altan-Bonnet, Paul François

## Abstract

The immune system’s most basic task is to decide what is “self” and “non-self”, but a precise definition of self versus non-self remains challenging. According to the discontinuity theory of immunity, effector responses depend on how quickly an antigenic stimulus changes: rapid change triggers an immune response, whereas gradual change fosters tolerance. We present a model of adaptive immune dynamics including T cells, Tregs and cytokines that reproduces the hallmarks of the discontinuity theory. The model allows for sharp discrimination between acute and chronic infections based on the growth rate of the immune challenge, and vaccination-like acute dynamics upon presentation of a bolus of immune challenge. We further show that the model behavior only depends on a handful of testable assumptions that we map to geometric constraints in phase space. This suggests that the model properties are generic and robust across alternative mechanistic details. We also examine the impact of multiple concurrent immune challenges in this model, and demonstrate the occurrence of dynamical antagonism, wherein, in some parameter regimes, slow-growing challenges hinder acute responses to fast-growing ones, with further counter-intuitive behaviors for sequential co-infections. Together, these results place the discontinuity theory on firm mathematical footing and encourage further investigation of interferences of multi-agent immune challenges, from chronic viral co-infections to cancer immunoediting.

## 1. INTRODUCTION

The conventional view of the immune system posits that it is focused on one goal: distinguishing and removing foreign molecules and cells (usually called “non-self”) from the host body (usually called “self”) [1, 2]. Systems immunology has made great progress in elucidating the parameters and characteristics of such immune discrimination [3], but, surprisingly, a quantitative definition of self vs non-self remains elusive. For instance, while biochemical parameters such as the binding affinities of immunogenic peptides to T cell receptors allow us to define effective “antigenicities” [4], their mapping to peptide sequences remains unclear. Statistical analysis of TCR receptors’ change during thymic selection revealed only small differences between positively and negatively selected sequences [5], and indeed self and non-self appear to have almost indistinguishable distributions in sequence space [6]. This suggests that unknown parameters besides sequence might play crucial roles in defining a proper immune response. Case in point: recent works have established that the immune environment in a broad sense plays a crucial role in modulating immune responses, e.g. due to cytokines and other costimulatory signals [7] or to antagonistic ligands that cancel the response to normally immunogenic peptides [8, 9].

There are multiple lines of evidence that the self/non-self dichotomy is not absolute. Our interior body is a complex dynamical system, itself interacting with a constantly changing environment. It thus makes sense that over very long time scales, our immune system adapts to these changes [10, 11]. This explains for instance why one can slowly be desensitized to allergens or why – when the process fails – auto-immune disorders appear, with (stochastic) flares of immune responses [12]. Understanding such adaptation is also of practical importance in treatments, e.g. to induce artificial tolerance to transplants [13] or in cancer immunotherapy. We know that multiple levels are implicated: for instance, it is well known that macrophages or NK cells can become tolerant to new antigens [14]. T cell responses can be primed and modulated by earlier exposure [8]. More generally, many systems-level feedbacks implicate specific tolerance genes [15] or Tregs [13]. Hence, what is recognized or not by the immune system might rather be a (slow) moving target, where immunogenic/nonimmunogenic categories do not perfectly align with self/non-self [16, 17]. To account for this feature, Pradeu, Vivier, and co-workers have proposed an alternative framework [18–22], illustrated in Fig. 1. As a first approximation, they propose that slow or gradual changes of immune challenges should be recognized as nonimmunogenic, allowing for the emergence of (new) immune tolerance with time (Fig. 1C). Conversely, any biochemical signal that changes rapidly in the organism is likely associated with a growing immune challenge, so should be immunogenic and lead to an acute response (Fig. 1B). Pradeu *et al*. thus proposed that the speed of change of molecular motifs with which immune cells interact is a determinant of immunogenicity, coining the “discontinuity theory” of immunity [19].

**FIG. 1.**
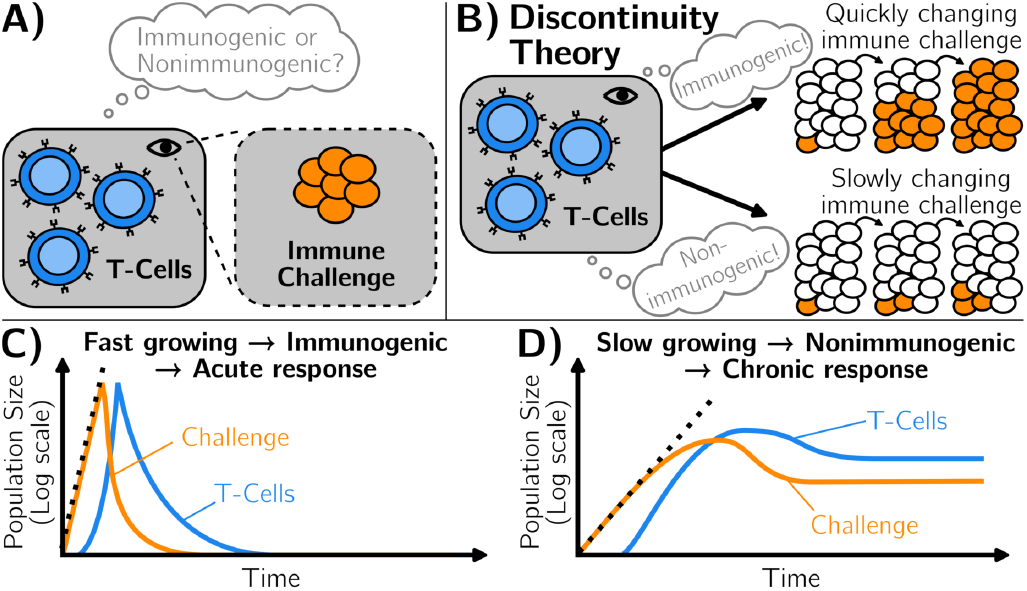
Illustration of the discontinuity theory of immunity. A) The immune system reacts to an immune challenge. It mounts a response that depends on its interaction with the challenge. B) The discontinuity theory of immunity hypothesizes that the immune system distinguishes between immunogenic and non-immunogenic challenges through the speed of change of its surroundings. C) Fast-growing challenges are perceived as immunogenic, triggering their elimination, through an acute immune response. D) Slow-growing challenges are perceived as non-immunogenic, which does not trigger their elimination, causing immune tolerance.

To date, an implementation of the discontinuity theory remains murky in regard to how immune discrimination can be determined by the speed of molecular change, both from the immunological and theoretical standpoints. Yet, there have been isolated observations in the field of quantitative immunology that suggest that biochemical derivatives, or at least time-dependent responses, may be critical in determining immunological outcomes (activation versus tolerance). For example, Mayer *et al*. explored how competition for a limited amount of antigen could decide the time range and overall extent of cell expansion [23]. Multiple groups also documented how the dynamic competition between effector and regulatory T cells (Tregs), based on the secretion and consumption of the cytokine IL-2, may decide immunological outcomes [10, 24– 27]. Reflecting this competition, a quantification of multiple cytokine dynamics showed how these signals universally encode an effective “immunological speed” determining longterm kinetics [4]. These models were classically analyzed in terms of quality and quantity of T cell activation deciding whether rapid/large IL-2 accumulation boosts effector functions, or, vice-versa, slow/meek IL-2 accumulation mostly boost the regulatory T cell compartment.

Here we revisit these ideas by introducing a coarse-grained version of the adaptive immune response – accounting for T cells, Tregs and cytokine response – and carrying out a thorough theoretical analysis to explore the discontinuity theory of immunology. We relate the discontinuity decision to the geometric properties of the dynamics in phase space, in particular to the *coexistence* of two types of trajectories corresponding to acute and chronic responses. We show that such coexistence strongly constrains the possible dynamics of the system, leading to specific bifurcations (in the dynamical systems theory sense) that naturally explain the extreme sensitivity to the rate of change of the immune challenge. To further establish this point, we leverage a new “landscape” framework [28] to build a purely geometric, “interaction-free” model of the system, recapitulating the properties of more explicit models. This approach proves that many properties of the description we propose are generic, i.e. expected in a broad family of models recapitulating the discontinuity theory of immunity, irrespective of their details.

Lastly, we study antagonism between immune challenges, whereby slow-growing challenges induce tolerance to fast-growing ones. Such effects are of primary importance for cases where immunological damages are close to self (e.g. tumors) or clinical trials of post-exposure vaccination against latent infections (e.g. mycobacterium tuberculosis, varicella-zoster etc.) [29, 30]. This study also provides a dynamical generalization of an immune spandrel theorem [31], specifying that absolute discrimination with respect to one kinetic parameter necessarily implies antagonism. Together, our results confirm that a modelling approach based on the description of simple dynamical rules (here, the co-existence of acute and chronic immune trajectories in response to growth rate) can lead to the derivation of non-trivial geometric properties, with clear experimental predictions for dynamical systems immunology.

## II. RESULTS

### A. A simple, minimal model recapitulates properties of the discontinuity theory of immunity

We start with an explicit model characterizing the dynamics of the adaptive immune system in response to an immune challenge (e.g. viral infection of cells, or growth of tumors), growing exponentially at the rate *r*. Infected cells linearly increase the activation and growth of effector T cells (called *T*), which in turn kill the infected cells *I*. Effector T cells exponentially grow in response to *I* and secrete the cytokine IL-2. IL-2, in turn, enhances the proliferation of effector T cells and regulatory T cells that mostly act as a sink for IL-2 in our simplified model. We further assume that, below some threshold, the immune challenge can not survive (see mathematical details in Supplement H), e.g. because there can be no fewer than 1 infected cell or because cooperative effects are necessary for an immune challenge to persist [32]. The model is illustrated in Fig. 2A, with equations given in Materials and Methods (Eqs 2). We chose parameters and interactions to approximate dynamics typically observed during an immune response (Table I), but we mostly use this model as an entry point to explore dynamical aspects and fundamental geometric properties of the immune response.

**TABLE 1.** Parameters used in the full model of 2 with their biological signification and value used in simulations. These are the values used for all figures, unless otherwise indicated. The time unit is “day”, i.e. all rates are implicitly per day.

| Symbol | Biological signification | Value |
| --- | --- | --- |
| $g$ | Linear growth rate | $10^1$ |
| $r$ | Exponential proliferation rate | 0.2 |
| $\kappa$ | Binding rate of T-cells to infected cells | $10^{-4}$ |
| $T^*$ | Concentration of naive T-cells at equilibrium | $10^2$ |
| $T_{\text{reg}}^*$ | Concentration of regulatory T cells at equilibrium | $10^4$ |
| $\alpha$ | T-cell proliferation rate | $10^4$ |
| $\beta$ | Consumption rate of IL2 by regulatory T-cells | 10 |
| $\delta_1$ | Inhibition rate of T-cells by regulatory T-cells | $10^{-3}$ |
| $\delta_2$ | Self-regulation rate of regulatory T-cells | $10^{-3}$ |
| $\gamma$ | Exhaustion rate of activated T-cells | 5.0 |
| $\mu$ | Production rate of IL2 by activated T-cells | $10^5$ |

**FIG. 2.**
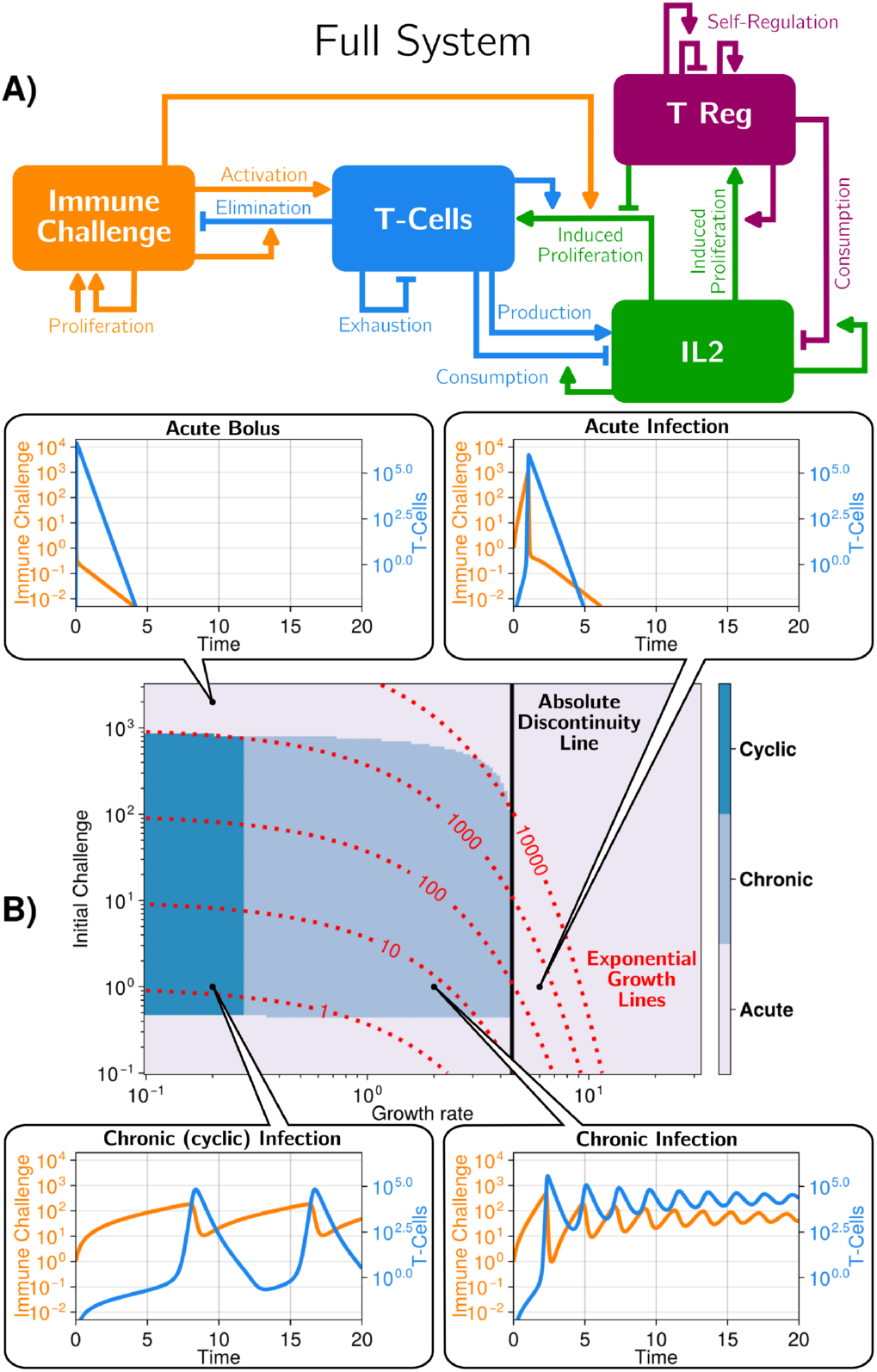
A model recapitulating the discontinuity theory of immunity A) Scheme of the model describing the interactions between an immune challenge, effector/regulatory T cells and cytokine (IL-2) in the adaptive immune system. Each arrow corresponds to an interaction in the model (e.g. the immune challenge activating the T-cells), full equations for the model are in the methods section, Eq. 2 with parameters in Table I. B) Different regimes in the parameter space defining the immune challenge (initial quantity of challenge, growth rate) for the full system. We show four typical trajectories. The system exhibits discontinuity in response to the growth rate, as illustrated by the vertical black line. For comparison, we add lines corresponding to a simple threshold model where immune response would be triggered only once a threshold of immune challenge is reached, see Supplement C and A for more details. Initial conditions: [T-Cells]= 0, [Tregs]= 10^4^, [IL2]= 0.

We define an immune response as acute when infected cells are eliminated (typically following an exponential increase in T cell numbers, followed by a decrease in both immune challenges and T cells), Fig. 1B. Conversely, following the discontinuity theory, for some other parameters, a regime might be reached where the immune challenge is not fully eliminated, coexisting with some (low-level) immune response: this situation defines a chronic response, Fig. 1C.

Fig. 2B illustrates the dynamics of the model for several values of the parameters defining the immune challenge, i.e. its growth rate *r* and initial value *I*(0) (other parameters we use are given in Table I). For graphical representation, we focus on the dynamics of the immune challenge and effector T cells (see Supplement, Fig. 7 for the full dynamics of the other variables). The model presents a broad variety of possible immune responses. For a high value of *r*, i.e. a fast-growing immune challenge, the immune response is always acute: both immune challenge and T cells grow exponentially fast (with an increase in IL-2 and of activated Tregs) (Fig. 2B, top). In this situation, the infected cells are quickly eliminated and the number of effector T cells slowly decreases until it becomes zero; meanwhile, both IL-2 and Treg number relax to their initial values. For lower *r* and low value of the initial immune challenge *I*(0), the dynamics are initially qualitatively similar, with a slower exponential increase in the number of both infected cells and effector T cells. The number of regulatory T cells and concentration of IL-2 also increase, but much less rapidly. Then, after some decrease and damped oscillations, all variables reach a non-zero, chronic equilibrium, characterized by a balance between infection of cells and effector T cell response (Fig. 2B, bottom). For even lower values of *r*, for our initial parameter choices, we observe another chronic regime where both T cells and infected cells oscillate with time.

The phase diagram in Fig. 2B illustrates the extent of chronic versus acute responses. Strikingly, there is a sharp boundary between chronic and acute regimes, around *r* = 4, with a vertical slope. This indicates that the nature of the response is based on the parameter *r only*. We contrast such a boundary with a hypothetical process where an immune response is triggered only when the immune challenge passes a pre-defined threshold after a fixed time (red dotted lines in Fig. 2B, see details in Supplement, section C).

Hence, our simple yet realistic model of immune response implements the main tenant of discontinuity theory: the rate of change in the immune challenge, *r*, decides whether effector T cells establish either an acute or a chronic response, irrespective of the precise initial immune challenge. We notice, however, that the chronic regime can only be reached for a range of initial challenge *I*(0). In particular, for high enough initial challenge *I*(0), the response is always acute, irrespective of *r* (“Acute bolus” in Fig. 2B). This is not inconsistent with discontinuity theory: mathematically, putting a high level of initial challenge at *t* = 0 is similar to having an extremely fast growing immune challenge challenge from the onset, so that we would indeed expect the immune system to yield an acute reaction if *I*(0) is high enough.

### B. Discontinuity theory : coexistence and disappearances of immune trajectories through bifurcations

To better understand the difference between chronic and acute regimes, we further reduce our model to two variables, *I* and *T*, by performing quasi-static approximations for [IL-2] and the number of Treg cells [33] (see details in Supplement F). This reduced model still displays similar properties of absolute sensitivity to the growth rate *r*, corresponding to the discontinuity theory (see comparisons in Supplement B), Fig. 3A. We derive an even simpler model, taking asymptotic limits for Tregs and IL-2 and rescaling variables and time (see materials and methods) such that we consider the following reduced 2D system:

**FIG. 3.**
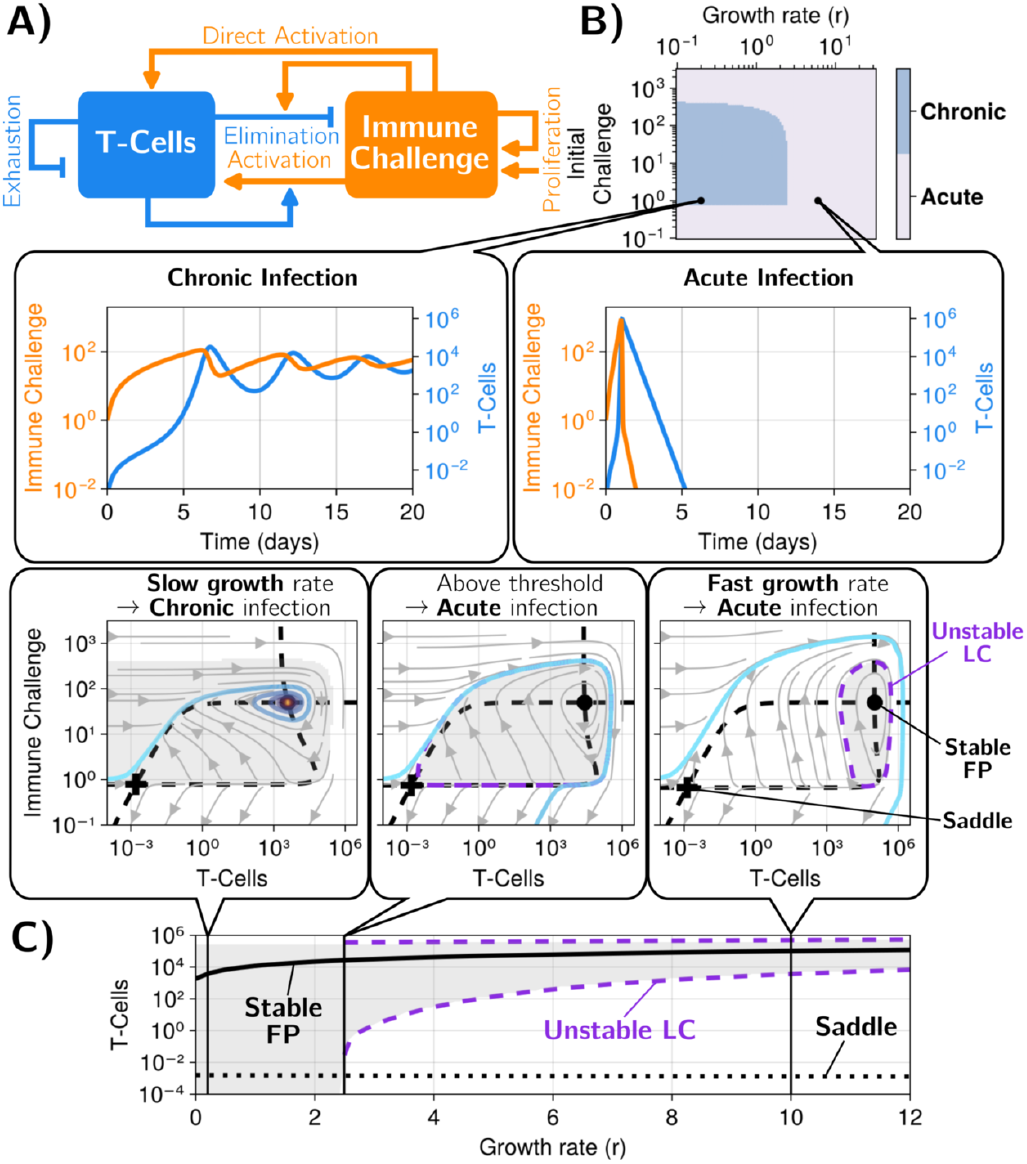
A) Network of interactions of the reduced system. This corresponds to equation 1. B) The different regimes in the parameter space of the reduced system. We show time series and phase portrait trajectories for the simulated systems. Examples of chronic (slow growth rate, pale blue region) and acute (fast growth rate, gray region) infections are presented. See supplement A for more details. C) Bifurcation diagram of the system as the growth rate (*r*) is varied. Snapshots of the phase portrait at different points along the bifurcation are shown. The shaded gray area corresponds to the basin of attraction of the chronic stable fixed point (FP). In the bifurcation diagram, we only shaded the region inside the unstable limit-cycle (LC). We use the same default parameters as in Fig. 2, Table I. Initial conditions: [T-Cells]= 0, [I]= 1.0

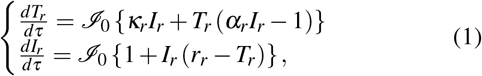

We explicitly introduce an operator, *I*_0_, indicating that the 2D field is interpolated towards the origin below the rescaled value *I*_0_, corresponding to the minimal quantity of immune challenge necessary to its survival (see Supplement H). *r*_*r*_ is the rescaled growth rate of the (rescaled) immune challenge, *I*_*r*_, which is depleted by (rescaled) T cells, *T*_*r*_. *I*_*r*_ in turn activates *T*_*r*_ with the (rescaled) rate *∀*_*r*_, and the growth rate of *T*_*r*_ itself depends on *I*_*r*_ (*#*_*r*_ term). Depletion/exhaustion of *T*_*r*_ occur with a rescaled rate of 1. This minimal 3-parameter model still displays exquisite sensitivity to the growth rate of the immune challenge, as illustrated in the phase diagram of Fig. 3B, top, while only keeping the acute and chronic regimes, as shown in the bottom panel (we represent trajectories in logscale). To interpret this model and to compare it to biological observables, we rescale *T*_*r*_ and *I*_*r*_ back to *T* and *I* appropriately in all relevant figures and discussion. The *r*_*r*_ parameter is also rescaled to *r* for easier comparison.

This reduced 3-parameter dynamical model, Eq. 1, combines features of different existing models. Its basic dynamics are close to previously proposed models of immune responses [34, 35] but it assumes additional nonlinearities for T cell growth. Those nonlinearities render the model more similar to generalized Lotka-Volterra models [36] with additional linear rates *∀*_*r*_*I*_*r*_ and 1 in equation 1. If we integrate the model defined in Eq. 1 without the interpolation *I*_0_ towards an attractive origin, we get damped solutions, similar to chronic ones, irrespective of the value of *r*_*r*_ (see Supplementary Figure 16 and Appendix section I). This points towards an important role of the nonlinear dynamics of immune challenge elimination when few of them are left.

Indeed, such 2D reduction of the dynamics allows us to perform phase-plane analysis in the *T, I* plane Fig. 3C. First, we observe that for all values of *r* (with scaling 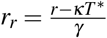), there exists a stable fixed point (solid black point in Fig. 3B-C bottom), which corresponds to the chronic state at lowest value of *r* (light blue trajectory in Fig. 3B-C, bottom left). Conversely, for very high *r*, the typical trajectory gets high in the *I, T* plane and circles back to the origin Fig. 3B, bottom right, as expected from an acute response. Crucially, this is true even if the initial challenge *I*(0) is very small (but non-zero). In other words, such acute response corresponds to an almost *homoclinic* trajectory [37] originating just above and circling back to the origin (light blue trajectory in 3B-C, bottom right). Notice that this acute trajectory circles around the stable fixed (solid black) point. We can then represent the basin of attraction of the fixed point (grey, Fig. 3C bottom middle and right), which is then limited by an *unstable limit cycle* separating the acute response trajectories from the (chronic) fixed point (dashed purple lines in Fig. 3C middle and right).

The unstable limit cycle is a direct mathematical consequence of the co-existence of a stable limit fixed point circled by a (almost) homoclinic trajectory. A homoclinic trajectory is an orbit in phase space starting from one point (technically a saddle point) and eventually converging to the same point after a long excursion. Such orbits are well known in neuroscience (e.g. neurons [38]). In an immune context, homoclinic trajectories are natural since, from a situation with no immune challenge, no T cells, a small injection of strong immune challenge should trigger an immune response, eventually going back to the initial (cleared) state. The existence of such dynamics is the fundamental reason explaining the extreme sensitivity in response to *r*. Indeed, as another consequence of the coexistence between an unstable limit cycle and a stable origin, there is a saddle point, close to the origin (black cross in Fig. 3C). As *r* decreases, the unstable limit cycle grows, until it collides with this saddle point close to 0, Fig. 3C bottom middle, and thus disappears (through a subcritical homoclinic orbit bifurcation [38], thus defining a true homoclinic orbit). Because of this bifurcation, acute trajectories such as the one in Fig. 3 C right, where low amount of initial challenges circle back to the origin, are no longer mathematically possible. Thus, one is left with the stable nonzero fixed point as the only stable attractor for many initial conditions *I*(0) in the absence of T cells above a threshold (gray zone represents the basin of attraction in Fig. 3C), ensuring chronicity for lower values of *r*.

So *r* acts as a control parameter for the bifurcation canceling out the unstable limit cycle, explaining the sharp transition between acute and chronic regime, and summarized in the bifurcation diagram at the top of Fig. 3C. Sensitivity to the growth rate only associated to discontinuity theory arises from the sudden change in the basin of attraction: the acute trajectories originating close to the origin for high *r* disappear, and are attracted to the chronic fixed point for low *r* for a broad range of *I*(0), compare Fig. 3C left with Fig. 3C right.

## III. LANDSCAPE GEOMETRIES FOR THE DISCONTINUITY THEORY OF IMMUNOLOGY

The properties described in the previous section suggest that the discrimination properties of the model are direct consequences of geometric features of the dynamical trajectories, in particular, the coexistence of acute and chronic trajectories in phase space. To confirm this intuition, we reverse engineer a minimal geometric model for discontinuity theory, using the Evoscape approach [28]. This approach, inspired by geometric modelling in biology [39] and kernel-based machine learning, relies on the combinations of simple dynamical modules to directly build “landscape-like” descriptions of biological systems in 2D, here the Immune challenge/effector T cell plane. We build landscapes based on two conditions, Fig. 4A :

**FIG. 4.**
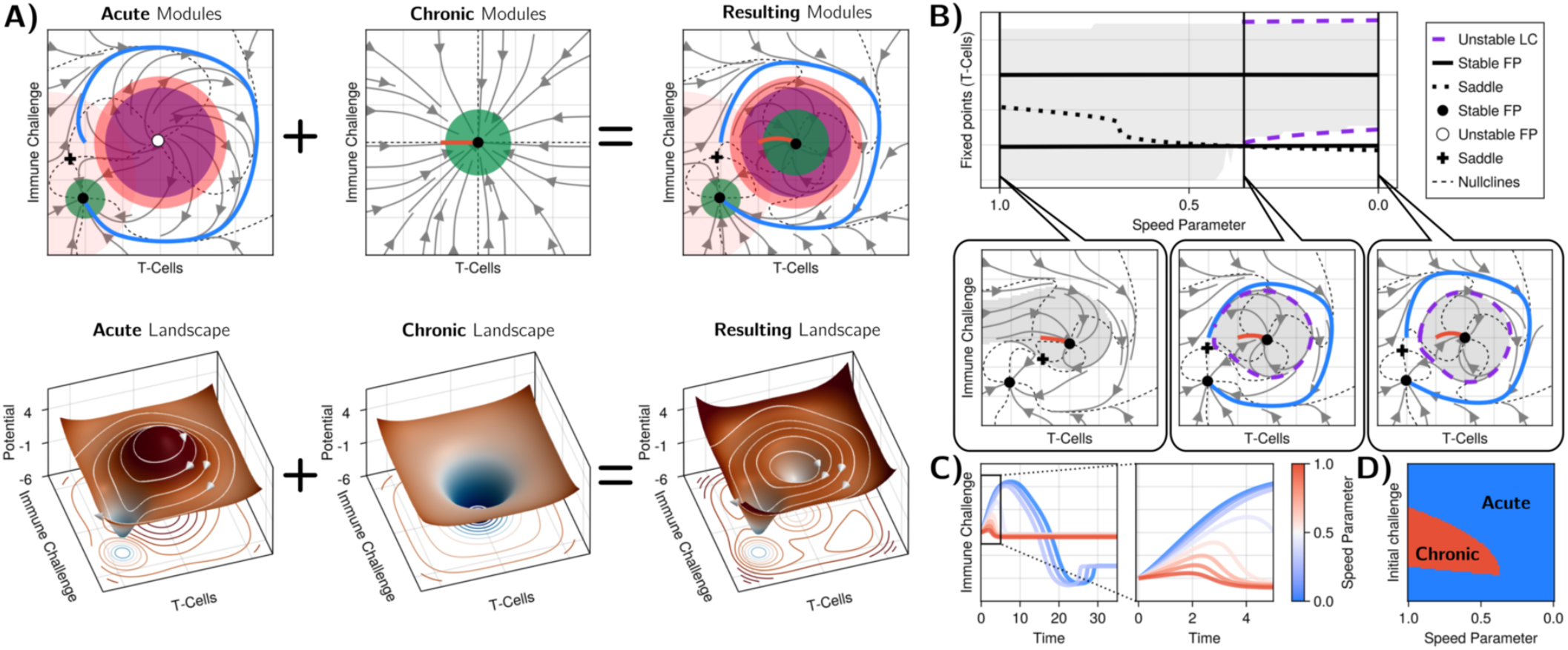
Reverse engineering a landscape model for the discontinuity theory of immunity A) Flows and potentials built using Evoscape[28]. The top row shows the flows, with the local Gaussian modules used indicated by circles. Green disks correspond to local attractors modules, purple rings to repelled, and orange modules to rotating flows. We also indicate examples of both acute and chronic trajectories. Landscapes corresponding to the flows are shown on the bottom row, with isolevels of the potential projected underneath the surface. The left column illustrates how to combine modules to get almost homoclinic trajectories corresponding to acute responses, circling to a bottom left attractor that would correspond to immune challenge elimination. The middle column is a simple attractor corresponding to the chronic response. Adding the two landscapes give a dynamic similar to what is observed in the explicit model, with acute/chronic trajectories going to two different attractors (green disks) separated by an unstable limit cycle. Equations can be found in Supplement B) Bifurcation diagram and phase plane analysis as the control (speed) parameter is varied, recapitulating the behaviours showing in Fig. 2C. C) Possible dynamics of the Immune Challenges as we vary the control parameter. D) Corresponding phase diagram.

1. there is an acute response for immune challenges achieving high growth rates *r*, even for very low initial values. This imposes a near-homoclinic acute trajectory from and to the origin in the limit of very high *r*, further associated to a stable fixed point at the origin. A corresponding landscape is shown in Fig. 4A, left.
2. there is a stable fixed point, topologically inside the acute homoclinic trajectory. A corresponding landscape is shown in Fig. 4A, middle

Equations for each landscape are given in Supplement.

The first condition is very natural for a functioning immune response: the systems should trigger an immune response even with a low number of immune challenges, then return to homeostasis after resolving the infection, analogous to an excitable-type trajectory [12, 28, 38]. Typically for such excitable/acute response, we expect the trajectory to circle an unstable attractor as seen in the Evoscape description in Fig. 4A, left. This first seems incompatible with the stability of a fixed point associated with the chronic response, Fig. 4A, middle.

Yet, one can build a simple phase space encompassing those two constraints simply using a linear combination of those two landscapes, Fig. 4A, right. As expected by design, we can get an almost homoclinic/excitable trajectory arising from the combination of a repeller with flow (acute trajectory, condition 1), itself around a stable fixed point in the center (chronic trajectory, condition 2). As a consequence, an unstable limit cycle naturally emerges. By varying the relative strength of modules (see details in Supplement), Fig 4B, the limit cycle can appear or disappear through a homoclinic bifurcation, and one can get either acute or chronic responses.

To fully match discontinuity theory, one needs to relate the control parameter to growth rate, which requires some extra condition. We thus chose to add an extra “growth” module close to the *y* axis for the “acute” landscape only (light orange disk in Fig. 4A left). The resulting model has all the hallmarks of discontinuity theory: fast-growing immune challenges reach the bottom attractor (corresponding to elimination), blue trajectories in Fig. 4C while slow-growing immune challenges reach the chronic attractor, red trajectories in Fig. 4C. One can then reconstruct a phase diagram delineating chronic from acute regimes, Fig. 4D, which is very similar to the ones derived from the models in Figs. 2-3B.

The fact that we can reproduce properties of the more explicit models using a reverse-engineered landscape suggest that the observed geometry and sequence of bifurcations are very natural, and should be shared by models with similar properties irrespective of their precise mathematical formulation, as long as they satisfy conditions 1-2 above.

## IV. DYNAMICAL ANTAGONISM AND INVERSION

The transition from acute to chronic regime, with a boundary controlled by a single parameter *r* is very reminiscent of specific and sensitive discrimination by T cells, which is sensitive to the kinetics of interaction of ligands to T cell receptor, almost irrespective of their concentration. Such behaviour has been well characterized experimentally [4, 8, 40, 41] and further studied theoretically [9, 42]. In particular, it was proved that such “absolute discrimination” mechanisms always display ligand antagonism [8, 43] as a phenotypic spandrel [44]. In those contexts, antagonistic properties occur at steady state. This motivated us to study antagonistic properties in the context of the discontinuity theory controlled by a dynamical bifurcation like here.

To do so, we expanded the simplified 3-parameter model of Eq. 1 to simulate a co-infection [45](we checked that the properties we describe below also hold for the expanded versions of the models). We consider two distinct immune challenges (*I*_1_ and *I*_2_) with different rates (*r*_1_ and *r*_2_) Fig. 5A-B. Dynamics of each of those immune challenges individually are presented in Fig. 5C-D. We assume that those challenges activate the same effector cells Fig. 5E. This situation is immunologically–realistic when immune challenges mutate to alter their infectivity and/or growth rate, while maintaining their antigenicity, which is now well established in multiple contexts from persistent bacteria [46] to tumor cells [47]. Equation 5 describes the dynamics of immune responses in these coinfections for the reduced system.

**FIG. 5.**
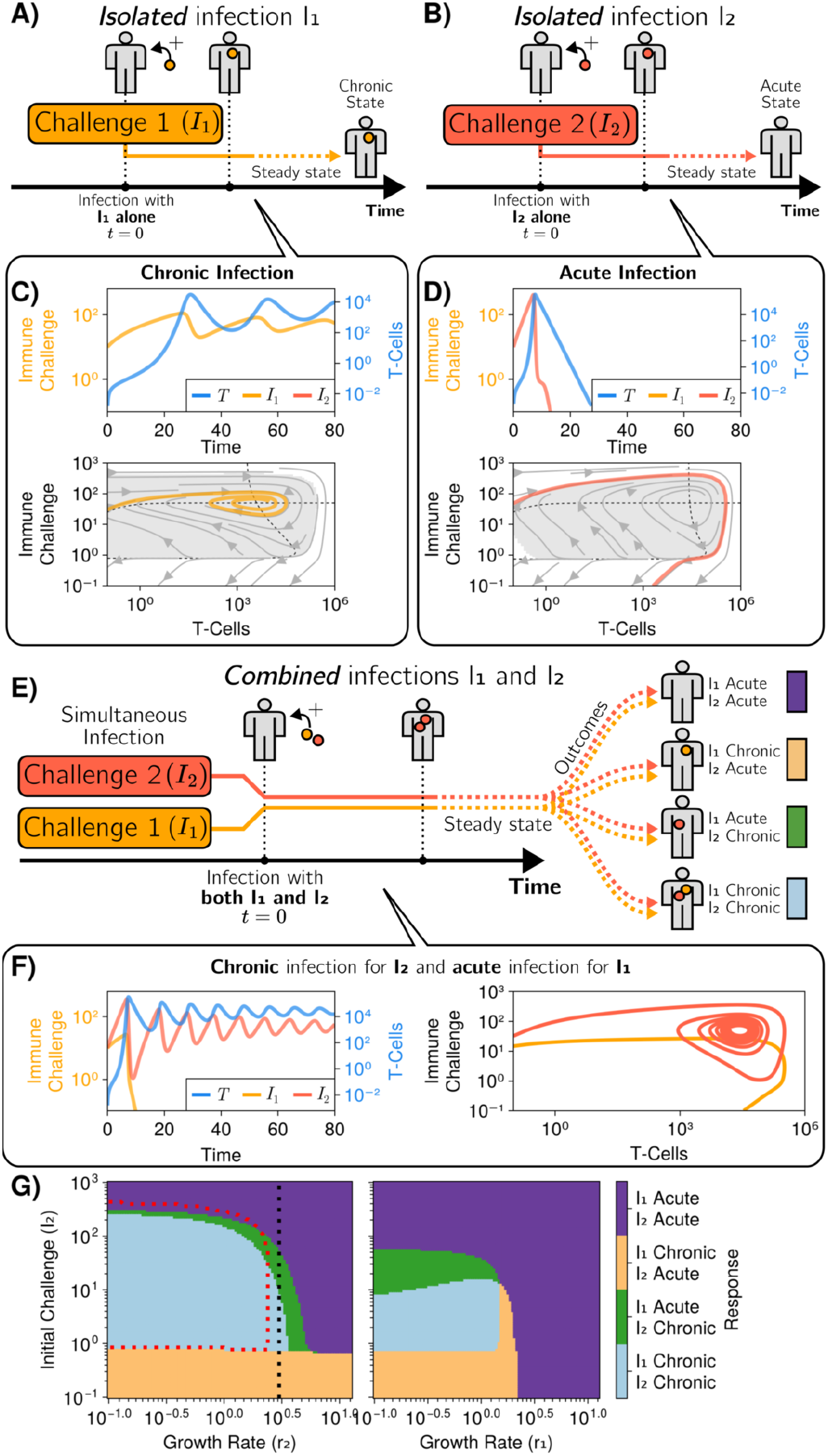
Dynamical antagonism and inversion A) Illustration of the dynamics for a single chronic response considered *I*_1_(0) = 10.0, *r*_1_ = 0.2 and B) for an acute response *I*_2_(0) = 10.0, *r*_2_ = 0.5, with phase plane dynamics and time courses displayed in C-D. Other parameters are the same as in Fig. 3E) Given the same two species presented in A and B, we can observe different outcomes to their isolated outcomes by combining them into a single infection (given by equation 5). F) Left: time series of two species combined, *I*_1_ and *I*_2_, undergoing, respectively, acute and chronic infections (the opposite of their original isolated outcomes, compare with dynamics in panels C-D). Right: trajectory in (*T, I*) space for each infection in the presence of the other one. G) Left: Infection outcomes for a combination of two species for different initial *I*_2_ and *r*_2_. The values of *I*_1_(0) and *r*_1_, leading to a chronic state for *I*_1_ when it is the only infectious species in the system, are fixed. The dotted red line represents the boundary between the chronic (inside) and acute (outside) regions when *I*_1_ is the only infectious species. Right: Crosssection of left plot, along the dotted black line. The growth rate for *I*_2_ (*r*_2_) is now fixed and the growth rate for *I*_1_ (*r*_1_) is allowed to move. *I*_1_(0)= 100.0, *T* (0)= 0.0

The dynamics of those co-infections both as a function of time and in phase space are shown in Fig. 5F. For this example, we observe a counter-intuitive inversion in the behaviour of both immune challenges Fig. 5F left : while the slow growing *r* challenge *I*_1_ is eliminated, the fast growing challenge *I*_2_ oscillates then stabilizes at the stable chronic fixed point. In particular, this means that a chronic immune challenge can antagonize an acute one, to give rise to a chronic regime for the fast-growing challenge (while the slow-growing challenge is eliminated).

Focusing on the phase plane trajectories, Fig. 5F right, we see that the trajectory of *I*_2_ is very similar to the acute trajectory, but converges to the chronic fixed point. To further understand what happens, in Supplement (Fig. 18D), we show the basin contours of the chronic fixed point for *I*_2_ as a function of the initial quantity of the chronically infected cells *I*_1_(0) : we see that addition of *I*_1_(0) modifies the unstable limit cycle, so that *I*_2_ crosses the boundary of the basin of attraction of the chronic fixed point very early during the simulated dynamics. As *I*_1_ increases, the basin contour retracts closer and closer to the chronic fixed point.

Those counter-intuitive effects depend on the respective values of growth rates and initial immune challenges, as shown in Fig. 5 G. We see four regimes, where each immune challenge can be either chronic or acute. The “inversion” regime corresponds to the green region where, when put together, a challenge *I*_1_ that would be chronic alone is becoming acute in co-infection, while a challenge *I*_2_ acute alone becomes chronic in co-infection. This regime thus occurs on the right of the normal chronic/acute boundary for *I*_2_ (dotted red line in 5 G). We also observe a regime where both *I*_1_ and *I*_2_ are chronic although *I*_2_ alone would be acute, which also extends on the right of the chronic/acute boundary for *I*_2_, again indicating antagonism on *I*_2_ by *I*_1_.

Because antagonism here is associated with the dynamics of immune challenges, there are in fact multiple possibilities depending on the timing when an acute or chronic challenge is added. We also consider the opposite limit, where one (chronic) challenge is already established, and another one is added later, Fig. 6A. This corresponds to a two-tier infection, sequentially stimulating the same immune cells, which could happen for instance if a mutation occurs for an immune challenge stabilized in the chronic regime.

**FIG. 6.**
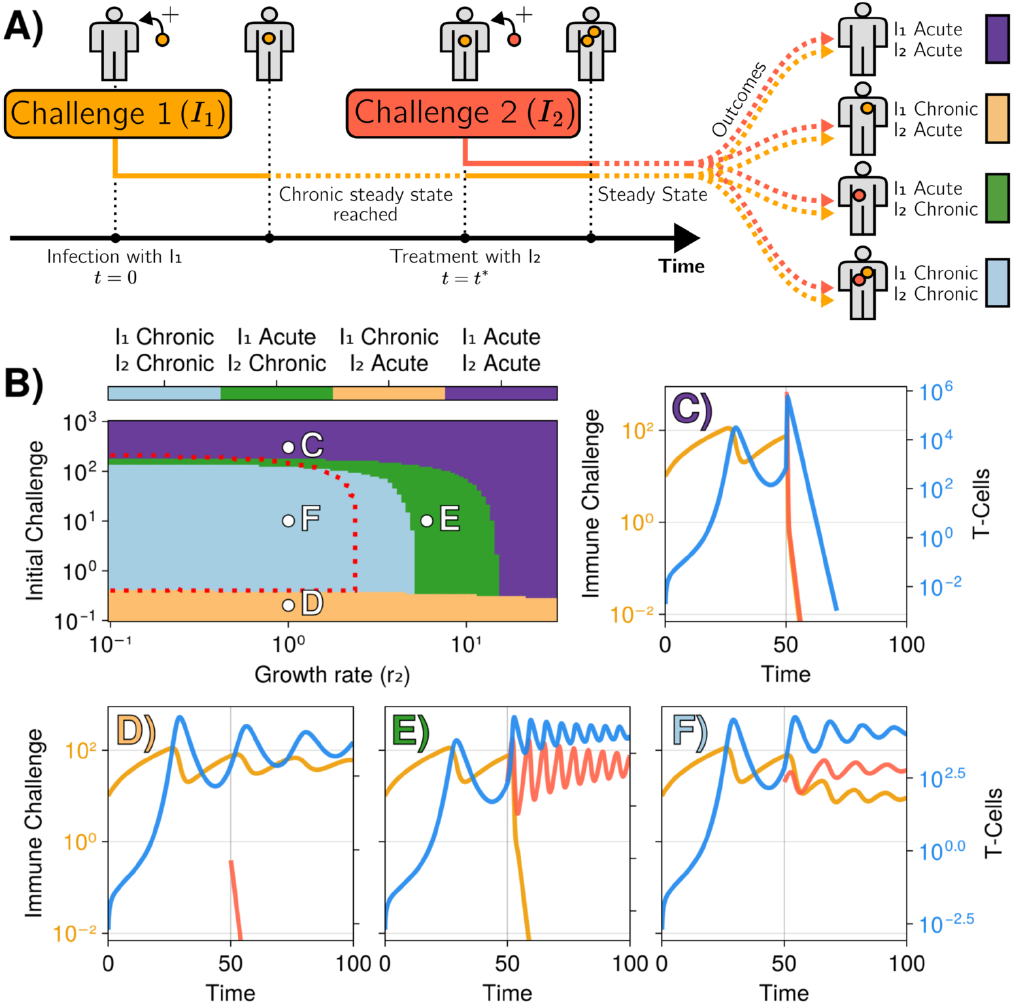
Illustration of the process of initial infection and subsequent “treatment”. A) The first immune challenge initially causes a chronic infection that reaches a steady state with *I*_1_ ≠ 0. Once this state is reached, a second immune challenge (*I*_2_) is added. Different outcomes can occur where each species can be either completely eliminated (acute) or remain active (chronic). Each case maps to cases presented in B-F. B) Infection outcomes for a system with initially a single species with *r*_1_ = 0.2. It evolves towards a chronic state. Once in this state, a second species is added, with its own *r*_2_, in amount *I* (*t*_treatment_). Four possible outcomes are possible and are color-coded. Examples of these processes and their outcomes are presented in subplots C,D,E,F. Parameters are similar to Fig. 3. *I*_1_(0)= 10.0, *T* (0)= 0.0

Fig. 6B-F illustrates the behavior of our 3-parameter model for this staggered co-infection, depending on the quantity and growth rate of the added immune challenge parameters *I*_2_(0), *r*_2_ (see e.g. [48] for an example of such coinfection mixing helminth and bacteria). Compared to Fig. 5 when both challenges are added initially, we see on the phase diagram Fig. 6B that both the antagonistic (light blue) and inversion (green) regions largely extend towards higher *r*, over more than half an order of magnitude in *r*. This means that the slow-growing immune challenge *I*_1_ in this region would typically drive a fast-growing one even more strongly towards the chronic regime. It is worth pointing out two other biologically relevant regimes. Panel E illustrates the inversion dynamics : while the initially chronic infection *I*_1_ is eliminated, the acute infection *I*_2_ is becoming chronic. Since *I*_2_ is growing about ten times faster than *I*_1_, this indicates a strong worsening of the chronic infection. Conversely, in panel C, if enough *I*_2_ is suddenly added, one can exit the chronic regime so that both challenges are eliminated. This is akin to a post-infection “vaccination” against the chronically established immune challenge *I*_1_.

## V. DISCUSSION

The discontinuity theory of immunity, proposed by Pradeu and collaborators, posits that the immune system is sensitive to the (quantitative) “speed of change” of immune challenges, rather than more qualitative features such as molecular signatures. It was used as a holistic framework to explain phenomena such as immune tolerance or autoimmune disease. Starting from a simple model of an immune response, displaying either acute/chronic activity depending on the growth rate *r* of the immune challenge, we derived a 3-parameter bidimensional model, recapitulating properties predicted by the discontinuity theory. In particular, the transition line separating acute and chronic regimes is purely controlled by the growth rate *r* over a broad range of initial sizes of immune challenges.

Our model presents common features to previously proposed models, for instance the co-existence of a chronic (“persistence”) state with a clearance state was proposed in [49], or an excitable model for auto-immunity proposed in [12]. However, those models did not consider the discontinuity theory framework. Our model presents unique, generic geometry, with excitable trajectories going *around* the persistence state in phase space. Such geometric constraints ensure that an unstable limit cycle separates the acute and chronic trajectories, and as a consequence the discontinuity detection occurs through a subcritical homoclinic orbit bifurcation depending on the growth rate of immune challenge *r*, canceling the unstable limit cycle. The properties described above would not depend on the particular details of the models, as long as the orbits can be reduced to 2D and a parameter such as *r* controls the acute to chronic regime. In particular, immunologists have documented with very high degree of granularity, how diverse immune responses take place depending on the type and size of infections (so-called Th1, Th2, Th17 etc. regimes for T cell responses).

The generic aspect is only expected close to the bifurcation when the acute and chronic trajectories co-exist. We can not exclude that different geometries and bifurcations are observed for more complex dynamics, e.g. effectively living in much higher dimensions. However, we notice that, if the transition from chronic to acute regime comes with a change of topology of the orbits, then by definition a global bifurcation is expected, and if acute immune responses indeed correspond to excitable dynamics [38], homoclinic bifurcations are a very natural scenario.

Antagonism is a generic property of multiple decisionmaking or discrimination pathways, from multiple immune recognition processes to olfaction [43, 50, 51], leading to practical application for cancer immunotherapy [52]. In [31], it was mathematically demonstrated that absolute discrimination, defined as ligand-based cellular decision-making based on one kinetic parameter irrespective of ligand concentrations, is necessarily associated with ligand antagonism, explaining why it is observed for many immune decisions [8]. We demonstrated here that discontinuity in the decision-making between acute and chronic response also displays antagonism, thus generalizing the results from [31] to a decision-making process based on a completely different mechanism (mathematical speaking), namely a global bifurcation.

This leads to important biological predictions. In a disease context, it raises the possibility of dynamical “adversarial” strategies [53] for a pathogen or a tumor to leverage dynamical antagonism to escape immune responses. For instance, a slow-growing tumor (like *I*_1_ in Fig. 6F) could first stabilize in the chronic regime of immunological response then later mutate into a faster-growing one (like *I*_2_) while keeping the immunological response at bay within a chronic regime. One could even imagine a slow ramp-up of immune escape of such increasingly growing tumors so that a fast-growing tumor that would normally trigger an immune response could slowly evolve and remain “undetected” by the immune system. It is especially important to point out that in our model, slow growing challenges are not passive and actively *antagonize* the detection of faster growing ones. Many observations about the differential growth rates of primary tumors vs metastasis, cancer dormancy as well differential immune responses (“immune privilege”), are consistent with such scenario [54, 55]. Of note, it has been recently observed that upon CAR-T treatment, tumors with slow growth tend to better survive [56]. This is consistent with both discontinuity theory and the observation in other contexts that persistor cells tend to escape treatment because of their slowed metabolism [46]. Conversely, the counterintuitive effects due to the immune challenge cross interactions could be used to tailor better treatments. For vaccine designs, bolus delivery is the normal regime (top left, acute regime in Figure 2B) but it may be tolerizing and inducing a chronic immune response if the inital challenge is too small/not antigenic enough. Such effect could further be leveraged to rather induce tolerance.

More generally, the coexistence of acute and chronic trajectories for the same immune challenge provides a simple, dynamical mechanism for an immune system to learn over time, thus allowing for a dynamic redefinition of the self [22]. Those predictions of 1. active antagonism of growing immune challenges and 2. mechanism of immune tolerance learning, naturally come from the geometric constraints imposed by discontinuity theory that we introduce. Similar to the general properties of antagonism for T cell detection [31], they come from the natural extension of a theoretical proposal for single immune challenges to interactions between multiple ones, with clear actionable experimental predictions. Practically, the existence and properties of such learning in immune dynamics could be directly tested using specialized platforms such as Immunotron [4]. In particular, by carefully monitoring time courses of acute/chronic infections, one should be able to identify topological changes in responses associated to global bifurcations such as the one predicted here.

Our model is of course very simplified and neglects multiple other mechanisms (e.g. sensitivity to the binding kinetics, immune editing, inflammatory switches, long term memory [57]), that could add multiple dimensions. That said, it is striking that a first-principle model accounting for discontinuity theory naturally comes with a combination of high-level features such as coexistence of acute and chronic responses, associated antagonism and global bifurcations discriminating between regimes. This is in line with the recent realization in multiple biological contexts that geometric, low-dimensional models accurately describe complex biological dynamics, e.g. line attractors for neural decision making [58] or heteroclinic flips for cellular differentiation [39, 59]. Of note, global bifurcations, transients and ghost states [60, 61] have been suggested to play important role complex cellular computations, and our model for discontinuity theory suggests that similar phenomena might be at play in immune decision-making.

## VI. MATERIALS AND METHODS

### A. Initial model and its variations

The starting point for describing the system is

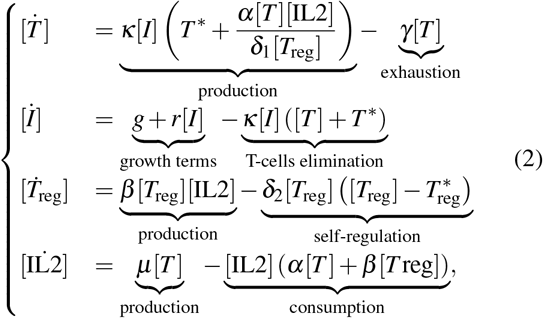

where [*I*] is the concentration of infected cells, [*T*] the concentration of T-cells, [*T*_reg_] the concentration of regulatory Tcells, and [IL2] the concentration of the interleukin 2, cytokine. The parameters used are described in table I. We complement those equations with an interpolating flow going towards the origin as soon as *I* is lower than a threshold *I*_0_ (see details in the Supplement H).

Assuming a quasistatic dynamic for [*T*_reg_] and [IL2], we can obtain a dynamical system of only two variables (see Appendix section F for derivation), which presents small differences with the full system (Fig. 10)

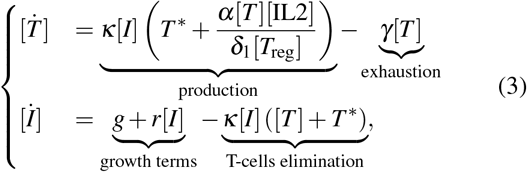

where [*T*_reg_] and [IL2] are functions of [*I*] and [*T*], given in equations F1 and F2. We designate this as the quasistatic system. We can do some further approximations and obtain and even more compact form:

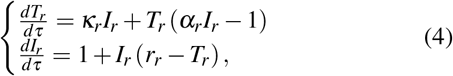

where the rescaling of the variables is given by 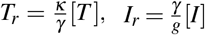, and 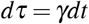 and the parameters are 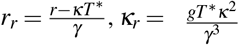 and 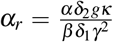. We also interpolate those equations towards the origin once *I*_*r*_ is lower than a threshold.

When describing the interactions of two species in the system like in section IV. We consider two species *I*_*r*,1_ and *I*_*r*,2_ that interact identically with the T-cells and do not interfere with each other directly. We describe the dynamics through

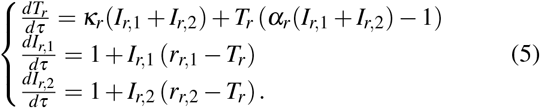

Notice that the immune challenges *I*_*r*,1_ and *I*_*r*,2_ have different growth rates, *r*_*r*,1_ and *r*_*r*,2_. These quantities can be rescaled to *r*_1_ and *r*_2_, the same way that *r*_*r*_ is rescaled to *r*. We can also simulate this two-species dynamics for the “full system” as well as the “quasistatic system” as shown in appendix G.

As stated in the main text, these reduced dynamical systems are rescaled back to their “natural” scale (from *I*_*r*_ and *T*_*r*_ to *I*_*r*_ and *T*_*r*_) when showing or discussing their simulated outcomes. The growth rate parameters *r*_*r*_, *r*_*r*,1_ and *r*_*r*,2_ are also rescaled to *r, r*_1_ and *r*_2_ for easier comparison with the full model, as well as allowing direct comparison with the biological observable that the parameters represent (the growth rate of the challenges).

### B. Dynamical landscapes

The dynamical landscape treatment presented in section III was conducted using the Evoscape framework [28]. The results of this analysis are all presented in Fig. 4.

Using 4 different types of modules (attractor, repellor, clockwise rotator and counter-clockwise rotator) with varying parameters we constructed the landscape presented in Fig. 4A. All the modules are linearly added together in a differential equation for I and T, with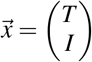,as such

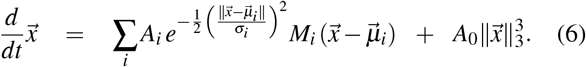

In the previous equation, all the modules are summed together, each denoted by an index *i*. Each of them have a location in phase space (the *T, I* plane) denoted by the vector *ωµ*_*i*_,a strength, denoted by *A*_*i*_, a width, denoted by *∋*_*i*_ and a Jacobian, denoted by *M*_*i*_. Furthermore, there is a global attractor with weight *A*_0_ (that we set to 0.01, for the combined landscape). This global attractor is of the form 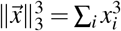. The Jacobian takes different form depending on the type of module. These are

- 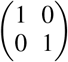: repellor
- 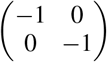: attractor
- 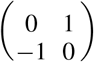: clockwise rotator
- 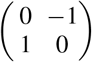: counter-clockwise rotator

Overall, the Evoscape framework allows us to mix and match different dynamical components to create a landscape presenting general features of the system. For the results of Fig. 4, we used the modules presented with their parameters in table II.

**TABLE 2.** Modules and their parameters used in the results of Fig. 4. Note that the strength of left rotator is dependent on the “Speed” parameter.

| Module Type | Strength | Width | Location |
| --- | --- | --- | --- |
| Attractor | 6.0 | 0.9 | $(-4, -3)$ |
| Repellor | 1.0 | 2.5 | $(0, 0)$ |
| Rotator (CW) | 0.6 | 3.0 | $(0, 0)$ |
| Rotator (CW) | Speed $\times 0.5$ | 4.0 | $(-6, -2)$ |
| Attractor | 3.3 | 1.5 | $(0, 0)$ |

The dynamics resulting from the Evoscape framework can be seperated into “potential” (attractors and repellors) and “curl” (rotators) parts. To draw the potentials presented in Fig. 4, we use the “potential” modules, from this, we can get an equation for the potential *P*:

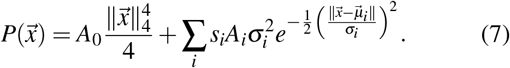

This is summing only over the repelling and attracting modules. *s*_*i*_ corresponds to the sign linked to each kinds of modules, such that *s*_*i*_ = −1 for an attractor (giving rise to a “valley” in the landscape) and *s*_*i*_ = + 1 for a repellor (giving rise to a “hill” in the landscape).

## ACKNOWLEDGMENTS

We thank Pankaj Mehta, Frédéric Guichard as well as the members of the François and Altan-Bonnet groups for useful discussions and comments. C.M. Denis is funded by the FRQ (DOI).

## REFERENCE

[1] R. Medzhitov and C. A. Janeway Jr, Decoding the patterns of self and nonself by the innate immune system, Science 296, 298 (2002).

[2] O. Feinerman, R. N. Germain, and G. Altan-Bonnet, Quantitative challenges in understanding ligand discrimination by αβ t cells, Molecular immunology 45, 619 (2008).

[3] M. Lever, P. K. Maini, P. A. Van Der Merwe, and O. Dushek, Phenotypic models of t cell activation, Nature Reviews Immunology 14, 619 (2014).

[4] S. R. Achar, F. X. P. Bourassa, T. J. Rademaker, A. Lee, T. Kondo, E. Salazar-Cavazos, J. S. Davies, N. Taylor, P. François, and G. Altan-Bonnet, Universal antigen encoding of T cell activation from high-dimensional cytokine dynamics, Science 376, 880 (2022), _eprint: https://www.science.org/doi/pdf/10.1126/science.abl5311.

[5] F. Camaglia, A. Ryvkin, E. Greenstein, S. Reich-Zeliger, B. Chain, T. Mora, A. M. Walczak, and N. Friedman, Quantifying changes in the t cell receptor repertoire during thymic development, Elife 12, e81622 (2023).

[6] A. Mayer, C. J. Russo, Q. Marcou, W. Bialek, and B. D. Greenbaum, How different are self and nonself?, arXiv 10.48550/arXiv.2212.12049 (2022).

[7] H. Yuan Kueh, A. Handel, A. Hoffmann, D. Chowell, R. A. Gottschalk, H. Singh, R. N. Germain, M. Meier-Schellersheim, K. Miller-Jensen, and G. Altan-Bonnet, What unique insights can modeling approaches capture about the immune system?, Cell Systems 15, 1148 (2024).

[8] G. Altan-Bonnet and R. N. Germain, Modeling t cell antigen discrimination based on feedback control of digital erk responses, PLOS Biology 3, e356 (2005).

[9] P. François, G. Voisinne, E. D. Siggia, G. Altan-Bonnet, and M. Vergassola, Phenotypic model for early T-cell activation displaying sensitivity, specificity, and antagonism, Proceedings of the National Academy of Sciences 110, E888 (2013).

[10] R. Marsland III, O. Howell, A. Mayer, and P. Mehta, Tregs self-organize into a computing ecosystem and implement a sophisticated optimization algorithm for mediating immune response, Proceedings of the National Academy of Sciences 118, e2011709118 (2021).

[11] T. Kato and T. J. Kobayashi, Understanding adaptive immune system as reinforcement learning, Physical Review Research 3, 013222 (2021).

[12] Y. Lebel, T. Milo, A. Bar, A. Mayo, and U. Alon, Excitable dynamics of flares and relapses in autoimmune diseases, Iscience 26 (2023).

[13] A. Cassano, R. Mora Cartin, P. Wang, Y. Wang, C. McIntosh, M. Andrade, A. Chong, and M.-L. Alegre, Role of tregs in maintaining alloreactive tconv hypofunction in transplantation tolerance, The Journal of Immunology 212, 1544–5560 (2024).

[14] B. N. Jaeger and E. Vivier, Natural killer cell tolerance: control by self or self-control?, Cold Spring Harbor Perspectives in Biology 4, a007229 (2012).

[15] M. Damo, N. I. Hornick, A. Venkat, I. William, K. Clulo, S. Venkatesan, J. He, E. Fagerberg, J. L. Loza, D. Kwok, et al., Pd-1 maintains cd8 t cell tolerance towards cutaneous neoantigens, Nature 619, 151 (2023).

[16] T. Pradeu and E. D. Carosella, On the definition of a criterion of immunogenicity, Proceedings of the National Academy of Sciences 103, 17858 (2006).

[17] T. Pradeu, Immunology and individuality, eLife 8, e47384 (2019).

[18] T. Pradeu, S. Jaeger, and E. Vivier, The speed of change: towards a discontinuity theory of immunity?, Nature Reviews Immunology 13, 764 (2013).

[19] T. Pradeu and E. Vivier, The discontinuity theory of immunity, Science immunology 1, aag0479 (2016).

[20] G. Eberl and T. Pradeu, Towards a general theory of immunity?, Trends in immunology 39, 261 (2018).

[21] T. Pradeu, The limits of the self: immunology and biological identity (Oxford University Press, 2011).

[22] T. Pradeu, Philosophy of biology, in The Philosophy of Science. A Companion (Oxford University Press; Oxford University Press, 2018).

[23] A. Mayer, Y. Zhang, A. S. Perelson, and N. S. Wingreen, Regulation of t cell expansion by antigen presentation dynamics, Proceedings of the National Academy of Sciences 116, 5914 (2019).

[24] S. Sakaguchi, N. Sakaguchi, M. Asano, M. Itoh, and M. Toda, Immunologic self-tolerance maintained by activated T cells expressing IL-2 receptor alpha-chains (CD25). Breakdown of a single mechanism of self-tolerance causes various autoimmune diseases., The Journal of Immunology 155, 1151 (1995).

[25] G. Voisinne, B. Nixon, A. Melbinger, G. Gasteiger, M. Vergassola, and G. Altan-Bonnet, T Cells Integrate Local and Global Cues to Discriminate between Structurally Similar Antigens, Cell Reports 11, 1208 (2015).

[26] H. S. Wong, K. Park, A. Gola, A. P. Baptista, C. H. Miller, D. Deep, M. Lou, L. F. Boyd, A. Y. Rudensky, P. A. Savage, et al., A local regulatory t cell feedback circuit maintains immune homeostasis by pruning self-activated t cells, Cell 184, 3981 (2021).

[27] S. Dikiy and A. Y. Rudensky, Principles of regulatory T cell function, Immunity 56, 240 (2023).

[28] V. Mochulska and P. François, Generative epigenetic landscapes map the topology and topography of cell fates, bioRxiv, 2025 (2025).

[29] H. Fu, J. A. Lewnard, I. Frost, R. Laxminarayan, and N. Arinaminpathy, Modelling the global burden of drug-resistant tuberculosis avertable by a post-exposure vaccine, Nature communications 12, 424 (2021).

[30] M. Ogilvie, Antiviral prophylaxis and treatment in chickenpox: a review prepared for the uk advisory group on chickenpox on behalf of the british society for the study of infection, Journal of Infection 36, 31 (1998).

[31] P. François, M. Hemery, K. A. Johnson, and L. N. Saunders, Phenotypic spandrel: absolute discrimination and ligand antagonism, Physical Biology 13, 066011 (2016).

[32] I. Andreu-Moreno and R. Sanjuán, Collective Infection of Cells by Viral Aggregates Promotes Early Viral Proliferation and Reveals a Cellular-Level Allee Effect, Current Biology 28, 3212 (2018).

[33] J. A. Borghans, R. J. De Boer, and L. A. Segel, Extending the quasi-steady state approximation by changing variables, Bulletin of mathematical biology 58, 43 (1996).

[34] H. Mayer, K. Zaenker, and U. An Der Heiden, A basic mathematical model of the immune response, Chaos: An Interdisciplinary Journal of Nonlinear Science 5, 155 (1995).

[35] E. D. Sontag, A dynamic model of immune responses to antigen presentation predicts different regions of tumor or pathogen elimination, Cell Systems 4, 231 (2017).

[36] W. Cui, R. Marsland III, and P. Mehta, Les houches lectures on community ecology: From niche theory to statistical mechanics, arXiv (2024).

[37] S. H. Strogatz, Nonlinear dynamics and chaos: with applications to physics, biology, chemistry, and engineering (Taylor and Francis, 2001).

[38] E. M. Izhikevich, Dynamical systems in neuroscience (MIT press, 2007).

[39] D. A. Rand, A. Raju, M. Sáez, F. Corson, and E. D. Siggia, Geometry of gene regulatory dynamics, Proceedings of the National Academy of Sciences 118, e2109729118 (2021).

[40] R. N. Germain and I. Stefanová, THE DYNAMICS OF T CELL RECEPTOR SIGNALING: Complex Orchestration and the Key Roles of Tempo and Cooperation, Annual Review of Immunology 17, 467 (1999).

[41] S. Farkona, E. P. Diamandis, and I. M. Blasutig, Cancer immunotherapy: the beginning of the end of cancer?, BMC Medicine 14, 73 (2016).

[42] J.-B. Lalanne and P. François, Principles of Adaptive Sorting Revealed by In Silico Evolution, Physical Review Letters 110, 218102 (2013).

[43] C. Torigoe, J. K. Inman, and H. Metzger, An unusual mechanism for ligand antagonism, Science 281, 568 (1998).

[44] P. François and G. Altan-Bonnet, The Case for Absolute Ligand Discrimination: Modeling Information Processing and Decision by Immune T Cells, Journal of Statistical Physics 162, 1130 (2016).

[45] S. Alizon and M. van Baalen, Multiple infections, immune dynamics, and the evolution of virulence, The American Naturalist 172, E150 (2008).

[46] R. A. Fisher, B. Gollan, and S. Helaine, Persistent bacterial infections and persister cells, Nature Reviews Microbiology 15, 453 (2017).

[47] Y. Goyal, G. T. Busch, M. Pillai, J. Li, R. H. Boe, E. I. Grody, M. Chelvanambi, I. P. Dardani, B. Emert, N. Bodkin, et al., Diverse clonal fates emerge upon drug treatment of homogeneous cancer cells, Nature 620, 651 (2023).

[48] K. Obieglo, X. Feng, V. P. Bollampalli, I. Dellacasa-Lindberg, C. Classon, M. Österblad, H. Helmby, J. P. Hewitson, R. M. Maizels, A. Gigliotti Rothfuchs, et al., Chronic gastrointestinal nematode infection mutes immune responses to mycobacterial infection distal to the gut, The Journal of Immunology 196, 2262 (2016).

[49] S. Baral, R. Antia, and N. M. Dixit, A dynamical motif comprising the interactions between antigens and cd8 t cells may underlie the outcomes of viral infections, Proceedings of the National Academy of Sciences 116, 17393 (2019).

[50] B. N. Dittel, R. N. Germain, C. A. Janeway, et al., Cross-antagonism of a t cell clone expressing two distinct t cell receptors, Immunity 11, 289 (1999).

[51] G. Reddy, J. D. Zak, M. Vergassola, and V. N. Murthy, Antagonism in olfactory receptor neurons and its implications for the perception of odor mixtures, Elife 7, e34958 (2018).

[52] T. Kondo, F. X. Bourassa, S. Achar, J. DuSold, P. F. Céspedes, M. Ando, A. Dwivedi, J. Moraly, C. Chien, S. Majdoul, A. L. Kenet, M. Wahlsten, A. Kvalvaag, E. Jenkins, S. P. Kim, C. M. Ade, Z. Yu, G. Gaud, M. Davila, P. Love, J. C. Yang, M. L. Dustin, G. Altan-Bonnet, P. François, and N. Taylor, Engineering TCR-controlled fuzzy logic into CAR T cells enhances therapeutic specificity, Cell 188, 10.1016/j.cell.2025.03.017 (2025).

[53] T. J. Rademaker, E. Bengio, and P. François, Attack and defense in cellular decision-making: lessons from machine learning, Physical Review X 9, 031012 (2019).

[54] J. A. Aguirre-Ghiso, Models, mechanisms and clinical evidence for cancer dormancy, Nature Reviews Cancer 7, 834 (2007).

[55] J. A. Joyce and D. T. Fearon, T cell exclusion, immune privilege, and the tumor microenvironment, Science 348, 74 (2015).

[56] A. L. Kenet, S. Achar, A. Dwivedi, J. Buckley, M. Pouzolles, H. Qin, C. Chien, N. Taylor, and G. Altan-Bonnet, The 1000+ mouse project: large-scale spatiotemporal parametrization and modeling of preclinical cancer immunotherapies, bioRxiv 10.1101/2025.03.17.643712 (2025), https://www.biorxiv.org/content/early/2025/03/20/2025.03.17.643712.full.pdf.

[57] M. Baliu-Piqué, M. W. Verheij, J. Drylewicz, L. Ravesloot, R. J. De Boer, A. Koets, K. Tesselaar, and J. A. Borghans, Short lifespans of memory t-cells in bone marrow, blood, and lymph nodes suggest that t-cell memory is maintained by continuous self-renewal of recirculating cells, Frontiers in immunology 9, 2054 (2018).

[58] M. Pagan, V. D. Tang, M. C. Aoi, J. W. Pillow, V. Mante, D. Sussillo, and C. D. Brody, Individual variability of neural computations underlying flexible decisions, Nature 639, 1 (2024).

[59] M. Sáez, J. Briscoe, and D. A. Rand, Dynamical landscapes of cell fate decisions, Interface focus 12, 20220002 (2022).

[60] D. Koch, A. Nandan, G. Ramesan, I. Tyukin, A. Gorban, and A. Koseska, Ghost channels and ghost cycles guiding long transients in dynamical systems, Physical Review Letters 133, 047202 (2024).

[61] L. Jutras-Dubé, E. El-Sherif, and P. François, Geometric models for robust encoding of dynamical information into embryonic patterns, Elife 9, e55778 (2020).

[62] J. Bezanson, A. Edelman, S. Karpinski, and V. B. Shah, Julia: A fresh approach to numerical computing, SIAM review 59, 65 (2017).

[63] C. Rackauckas and Q. Nie, DifferentialEquations.jl – A Performant and Feature-Rich Ecosystem for Solving Differential Equations in Julia, JORS 5, 15 (2017).

[64] D. P. Sanders, L. Benet, B. Richard, J. Grawitter, E. Gupta, L. Ferranti, D. Karrasch, O. Hénot, Z. Hurák, Y. Sharma, T. Frondelius, M. Forets, J. TagBot, G. Datseris, E. Schnetter, E. Hanson, and E. Saba, JuliaIntervals/IntervalRootFinding.jl: v0.6.0 (2024).

[65] S. Danisch and J. Krumbiegel, Makie.jl: Flexible high-performance data visualization for Julia, Journal of Open Source Software 6, 3349 (2021).

[66] A. M. Kramer, L. Berec, and J. M. Drake, Editorial: Allee effects in ecology and evolution, Journal of Animal Ecology 87, 7 (2018), _eprint: https://onlinelibrary.wiley.com/doi/pdf/10.1111/1365-2656.12777.

[67] G.-Q. Sun, Mathematical modeling of population dynamics with Allee effect, Nonlinear Dyn 85, 1 (2016).

