## supplementary information for "Dynamical model and geometric insights in the discontinuity theory of immunity"

### Appendix A: Numerical tools

All simulations were done using the Julia language [62]. The integration was carried out using the `DifferentialEquations.jl` [63]. Specifically, the `Rodas4P` integration scheme was used for all simulations. Fixed points where computed using the `IntervalRootFinding.jl` package [64].

The plots were created using the `Makie.jl` (CairoMakie) library [65].

To generate the phase diagram in figure 2B, we sorted out the trajectories once that they had reached a steady-state (we assumed that this was attained after around 160 days). This steady state was observed for around 40 days and a decision tree for the classification of the outcomes was made. In pseudo code, this was:

```
challenge = steadyStateTimePoints()

if (challenge[last] < 1)
    return Acute
else
    if (maximum(challenge) - minimum(challenge) < 1)
        return Cyclic
    else
        return Chronic
    end
end
```

The outcomes are validated by the example trajectories presented in figure 2B. For figure 3B, because of the knowledge of the absence of a stable limit cycle, we only numerically checked for the `Chronic` and `Acute` outcomes. We used the same previous procedure, but with fewer time steps (10 days) since, typically, acute trajectories rapidly circle the basin of attraction to reach an acute state. Again, this was also further verified through the plotted time-courses and phase-portraits.

The notebooks and code used to generate the simulations and figures can be accessed at this repository.

### Appendix B: Dynamics of IL2 and regulatory T-Cells

The dynamics of the IL2 and Tregs are coupled to those of the T-Cells which in turn is coupled to the number of infected cells as illustrated in Fig. 3A. Figure 7 shows how those quantities behave over time. This figure is to be compared with the time series in Fig. 3D-F).

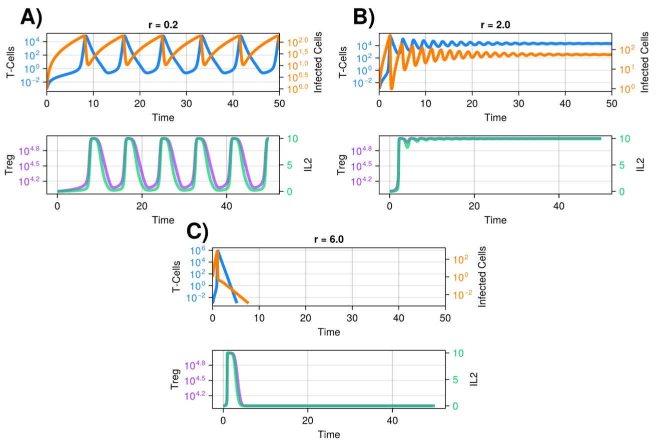


FIG. 7. Time series for the full system presented in Fig. 2. Here the courses for IL2 and the Tregs are included. Initial conditions: [T-Cells] = 0, [Immune Challenge] = 1.0, [Treg] = $10^4$, [IL2] = 0.

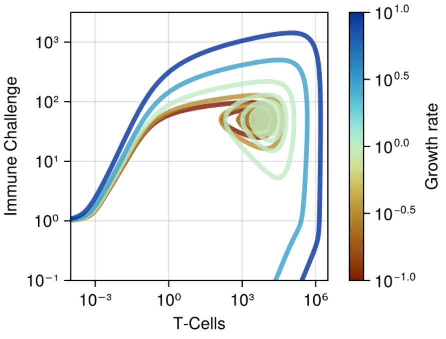


FIG. 8. Trajectories in the $I$-$T$ space for the reduced system for different proliferation rates. In the beginning of the infections, higher growth rates have steeper slopes than lower ones. Initial conditions: [T-Cells] = 0, [Immune Challenge] = 1.0.

### Appendix C: Exponential growth curves

In Fig. 3B, we plot "exponential growth curves". This is to show that the absolute discontinuity observed in the model is not merely a consequence of the exponential growth of the challenge. In fact, these contour lines are not as vertical as the boundary between the chronic and acute states.

These delineate how much of the challenge would be in the system after a time $\tau$ has passed, given a growth rate, $r$, and an initial quantity for this challenge. All the points along a growth line $N$ will have the challenge in the amount $N$ after time $\tau$. Mathematically, this means $N = I_0 e^{r\tau}$. Since every point in the phase diagram of Fig. 3B corresponds to a $I_0$ and a $r$, this allows us to associate a $N$ to each of them, and consequently draw "contour lines" for a given $N$.

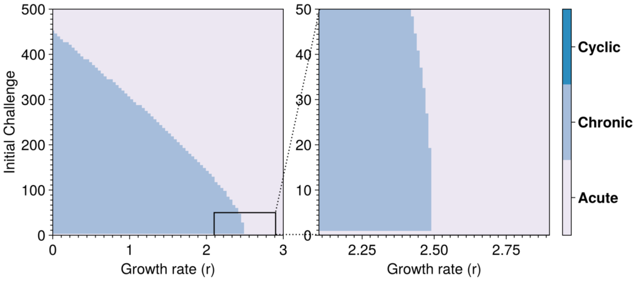


FIG. 9. Reproduction of the results presented in Fig. 3B on a linear scale instead of a logarithmic scale. The right plot zooms in on a region of the plot on the left.

#### Appendix D: Growth rate and steepness of trajectories

In Fig. 8 we vary the growth rate ($r$) and show the resulting trajectories. This highlights their change in slope. We see that the two trajectories with the highest growth rates are above the threshold and yield acute infections (they are not going towards the chronic fixed point like the other).

#### Appendix E: Linear scale plots

The plots in the main text have been presented on a logarithmic scale to see some details better. However, we can also produce them in linear scale as shown in Fig. 9.

#### Appendix F: Derivation of reduced model

Starting with equation 2, we approximate the variables $[T_{\text{reg}}]$ and [IL2] as quasistatic, i.e. $[\dot{T}_{\text{reg}}] = [\dot{\text{IL2}}] = 0$. This allows us to obtain analytic expressions for both of these in terms of $[T]$, $[I]$ and the parameters. At a given moment in time, given the number of T-cells $[T]$ and of infected cells $[I]$, we can entirely specify the number of regulatory T-cells $[T_{\text{reg}}]$ and of interleukin-2 [IL2] by

$$[T_{\text{reg}}] = \frac{-\delta_2\left(\alpha[T] - \beta T^*_{\text{reg}}\right) + \sqrt{\left([T]\alpha\delta_2^2 + T^*\text{reg}\beta\delta_2\right)^2}}{2\beta\delta_2} \tag{F1}$$

and

$$[\text{IL2}] = \frac{-\delta_2\left(\alpha[T] + \beta T^*_{\text{reg}}\right) + \sqrt{\left([T]\alpha\delta_2^2 + T^*\text{reg}\beta\delta_2\right)^2}}{2\beta^2}. \tag{F2}$$

Notice two symbolic differences in these: a sign difference in the first term of the numerator and the factor of the denominator. Using these forms for $[T_{\text{reg}}]$ and [IL2] in equation 2 gives us the quasistatic approximation of equation 3. Notice that we can also relate these last two quantities via $[\text{IL2}] = \frac{\delta_2}{\beta}\left([T_{\text{reg}}] + T^*_{\text{reg}}\right)$.

We can further reduce this system for large $[T]$, which makes the fraction $\frac{[\text{IL2}]}{[T_{\text{reg}}]}$ constant, and simplifies the dependence on $[T_{\text{reg}}]$ and [IL2]. We have

$$\lim_{[T]\to\infty} \frac{[\text{IL2}]}{[T_{\text{reg}}]} = \lim_{[T]\to\infty} \frac{\frac{\delta_2}{\beta}\left([T_{\text{reg}}] + T^*_{\text{reg}}\right)}{[T_{\text{reg}}]} = \frac{\delta_2}{\beta}.$$

Hence, we can now write the system as

$$\begin{cases} [\dot{T}] &= \underbrace{\kappa[I]\left(T^* + \frac{\alpha\delta_2}{\beta\delta_1}[T]\right)}_{\text{production}} - \underbrace{\gamma[T]}_{\text{exhaustion}} \\ [\dot{I}] &= \underbrace{g + r[I]}_{\text{growth terms}} - \underbrace{\kappa[I]\left([T] + T^*\right)}_{\text{T-cells elimination}}. \end{cases} \tag{F3}$$

Substituting $[T] = \frac{\gamma}{\kappa}T_r$, $[I] = \frac{g}{\gamma}I_r$ and $dt = \frac{1}{\gamma}d\tau$ and rearranging allows us to obtain equation 4 where we substituted the parameters $r_r = \frac{r-\kappa T^*}{\gamma}$, $\kappa_r = \frac{gT^*\kappa^2}{\gamma^3}$ and $\alpha_r = \frac{\alpha\delta_2 g\kappa}{\beta\delta_1\gamma^2}$.

We compare the numerical behavior of the quasistatic system and the full system in Fig. 10. An interesting characteristic of the structure of this system is its similarity with the Lotka-Volterra system. Figure 16A compares the network of interactions between the two systems. Furthermore, looking at the system without any kind of threshold (acting like an Allee effect mechanism [32]), we observe similar dynamics to Lotka-Volterra type systems in Fig. 16B-C.

#### Appendix G: Multi-challenge simulations for full and quasistatic system

The figures 14, 12, 15, 13 present the results of the multi-challenge simulations for the full system (equation 2) and the quasistatic system (equation 3) for the "combined scenario". This is to be compared with Fig. 5: both challenges are put at the same time in the system. We also show simulations for the "treatment scenario". This is to be compared with Fig. 6: the second challenge is added to the system after the first one has reached its chronic state. Both the full system and the quasistatic system present oscillations and consequently contribute to more possible infection outcomes than the reduced system (equation 4) e.g. $I_1$ is chronic and oscillating, $I_2$ is chronic without oscillations. However, notice that we do not observe all possible permutations of chronic, acute and oscillatory behavior in the multi-challenges simulations. The overall structure of these phase diagrams is nonetheless very similar to those of figures 5 and 6.

#### Appendix H: Threshold and Allee effect

A core assumption of our model is that below a certain quantity of infected cells, its number falls to zero. This is akin to the introduction of an Allee effect in ecology, where below a certain capacity, the population dynamics fall off to

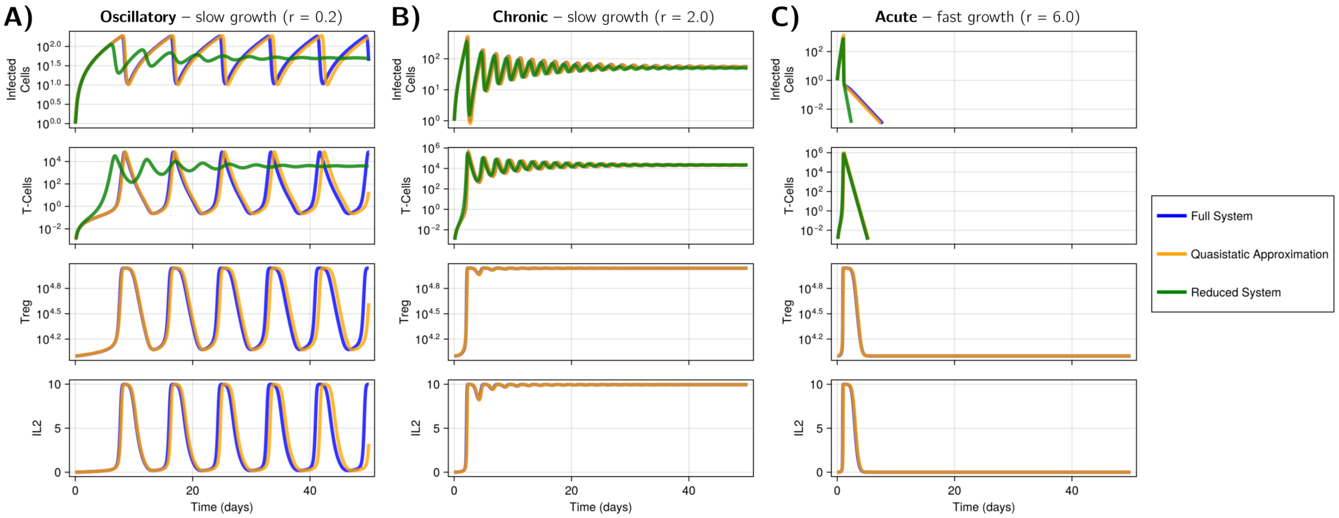


FIG. 10. Numerical solutions to equations 2, 3 and 4 for three different challenge growth rates (A, B and C). The quasistatic (equation 3) and reduced (equation 4) systems agree well with the full system (eq 2), except the reduced system, for small growth rate, which does not present limit cycle oscillations. Note that the reduced model only has trajectories for infected cells and T-cells, but none for the regulatory T-cells and IL2. Acute case: $r = 6.0$. Chronic case: $r = 2.0$. Chronic oscillatory case: $r = 0.2$. Initial conditions: $[\text{T-Cells}] = 0$, $[\text{Immune Challenge}] = 1.0$, $[\text{Tregs}] = 10^4$, $[\text{IL2}] = 0$.

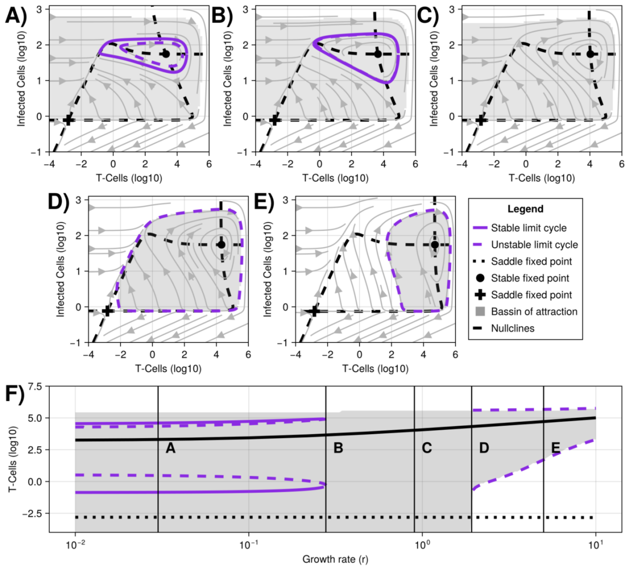


FIG. 11. A-E) Phase portraits for the quasistatic system (equation 3) for different $r$ parameters. Note that there is also a fixed point at the origin that is not shown due to the log-scale nature of the plots. A) $r = 0.03$, Coexistence of stable and unstable limit cycles circling a stable fixed point. B) $r = 0.278$, Saddle node (SN) bifurcation occurring between the two limit cycles. C) $r = 0.9$, Only a fixed point and its basin of attraction are left after the SN bifurcation. D-E) $r = 1.92434$ and $r = 5$, Homoclinic bifurcation transforms a basin of attraction into an unstable limit cycle with a stable fixed point at its center. F) Bifurcation diagram for the quasistatic system along the $r$ parameter axis. The dynamics occurring at the vertical bars are plotted in subplots A-E.

zero entirely for many possible reasons (e.g., too few individuals available for breeding, collective advantage when hunting not existent, etc.) [32, 66, 67]. We implemented two different kinds of thresholds in this work. First is a "hard threshold", where the number of infected cells falls off exponentially under a certain quantity (we fixed it at $I = 1$, in our case). This means that the equation for the number of infected cells is piecewise, as such:

$$\dot{[I]} = \begin{cases} g + r[I] - \kappa[I]\left([T] + T^*\right), & \textbf{if } [I] \geq 1 \\ -[I], & \textbf{if } [I] < 1. \end{cases} \tag{H1}$$

Second, a "soft threshold" was introduced and is used in all simulations of this work except where explicitly mentionned. We implement a continuous variation across the threshold between the "below capacity" and "above capacity" states via a hill function for this case. We hence use an expression for the infected cells:

$$\dot{[I]} = \left(g + r[I] - \kappa[I]\left([T] + T^*\right)\right) H_{n,k}\left([I]\right) - [I]\left(1 - H_{n,k}\left([I]\right)\right). \tag{H2}$$

Where $H_{n,k}\left([I]\right) = \frac{\left(\frac{[I]}{k}\right)^n}{\left(\frac{[I]}{k}\right)^n + 1}$. This Hill function acts a way to smoothly interpolate or "cross-fade" between the two pieces of equation H1. The shapes of this curve are presented in Fig. 17. The parameter $n$ controls how steep the interpolation is and the parameter $k$ controls where the threshold is located. For large $n$ we have a Heaviside function that is identical to the "hard threshold". Figure 16 shows how the phase portrait behaves when not taking into account this threshold.

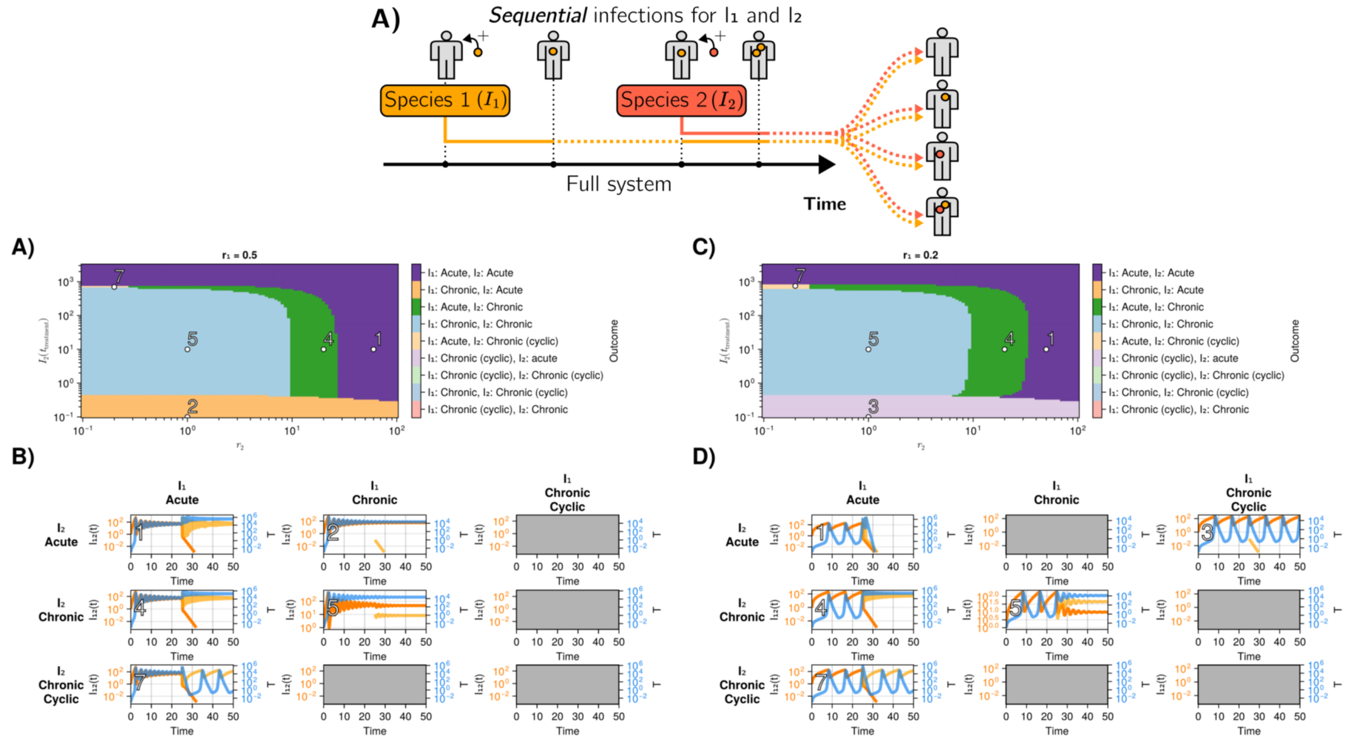


FIG. 12. A) Infection outcomes for the full system (equation 2) in the "treatment scenario", with initially a single species with $r_1 = 0.5$. It evolves towards a chronic state. The figure is to be compared with Fig. 6. At $t_{\text{treatment}} = 25$, a second species is added, with its own $r_2$, in amount $I\left(t_{\text{treatment}}\right)$. Examples of these processes are presented in sub-subplots B). B) Example time series associated with the labeled points in A). Some combinations are not attainable. C) Same as A), but the initial chronic state is in a cyclic state, with $r_1 = 0.2$. D) Example time series associated with the labeled points in C). Notice that some of the colors in the colorbar do not appear in the phase diagram.

We extended this threshold for the reduced system as well by taking into account the threshold point via the proper scaling of variables. In nondimensional units we write

$$\frac{dI_r}{d\tau} = \left(1 + I_r\left(r_r - T_r\right)\right) H_{n,k}(I_r) - I_r\left(1 - H_{n,k}(I_r)\right) \tag{H3}$$

### Appendix I: Fixed points of reduced model

For the reduced model, we can obtain and characterize the fixed points of the system. Starting with

$$\begin{cases} \frac{dT_r}{d\tau} = \kappa_r I_r + T_r\left(\alpha_r I_r - 1\right) \\ \frac{dI_r}{d\tau} = \left(1 + I_r\left(r_r - T_r\right)\right) H_{n,k}(I_r) - I_r\left(1 - H_{n,k}(I_r)\right), \end{cases} \tag{I1}$$

we set both equation to 0. This gives $T_r = \frac{\kappa_r I_r}{1-\alpha_r I_r}$. We can obtain the fixed points by solving the equation

$$0 = I_r^2\left(\left(\frac{I_r}{k}\right)^{n-2}\left((r_r\alpha_r - \kappa_r)I_r^2 + (\alpha_r - r_r)I_r - 1\right) - \alpha_r\right). \tag{I2}$$

However, practically, to compute the fixed points, in particular those appearing in streamplots, we use the `IntervalRootFinding.jl` package [64].

#### 1. Stability analysis in the absence of interpolation towards the origin

In the absence of interpolation towards the origin (i.e., $H_{n,k}(I_r) = 1$ in equation I1 above), we can analytically obtain the fixed point $(T_r^*, I_r^*)$ and its linear stability. First, solving for fixed points, we express $I_r^*$ in terms of $T_r^*$ and solve the resulting quadratic equation for $T_r^*$, finding

$$\begin{aligned} T_r^* &= \frac{r_r + \alpha_r}{2} + \frac{1}{2}\sqrt{(r_r + \alpha_r)^2 + \kappa_r} \\ I_r^* &= \frac{1}{T_r^* - r_r} = \frac{T_r^*}{\kappa_r + \alpha_r T_r^*}\,. \end{aligned} \tag{I3}$$

Only the $+$ root is in the quadrant of interest, since the square root $\sqrt{(r_r + \alpha_r)^2 + \kappa_r} > r_r + \alpha_r$ for any biologically meaningful (i.e., positive) value of $\kappa_r$. This also ensures that $T_r^* > r_r > 0$, such that $I_r^* > 0$ as well.

Second, we note that trajectories in the upper quadrant stay in the upper quadrant, since $\dot{T}_r(T_r = 0) = \kappa_r I_r > 0$ (since $I_r \geq 0$ in the upper quadrant), and $\dot{I}_r(I_r = 0) = 1 > 0$. Hence, the Poincaré-Bendixson theorem guarantees there must be an attractor in this quadrant (no chaos in 2D).

Third, we perform a linear stability analysis around $T_r^*, I_r^*$. The Jacobian matrix of the system, evaluated at the fixed

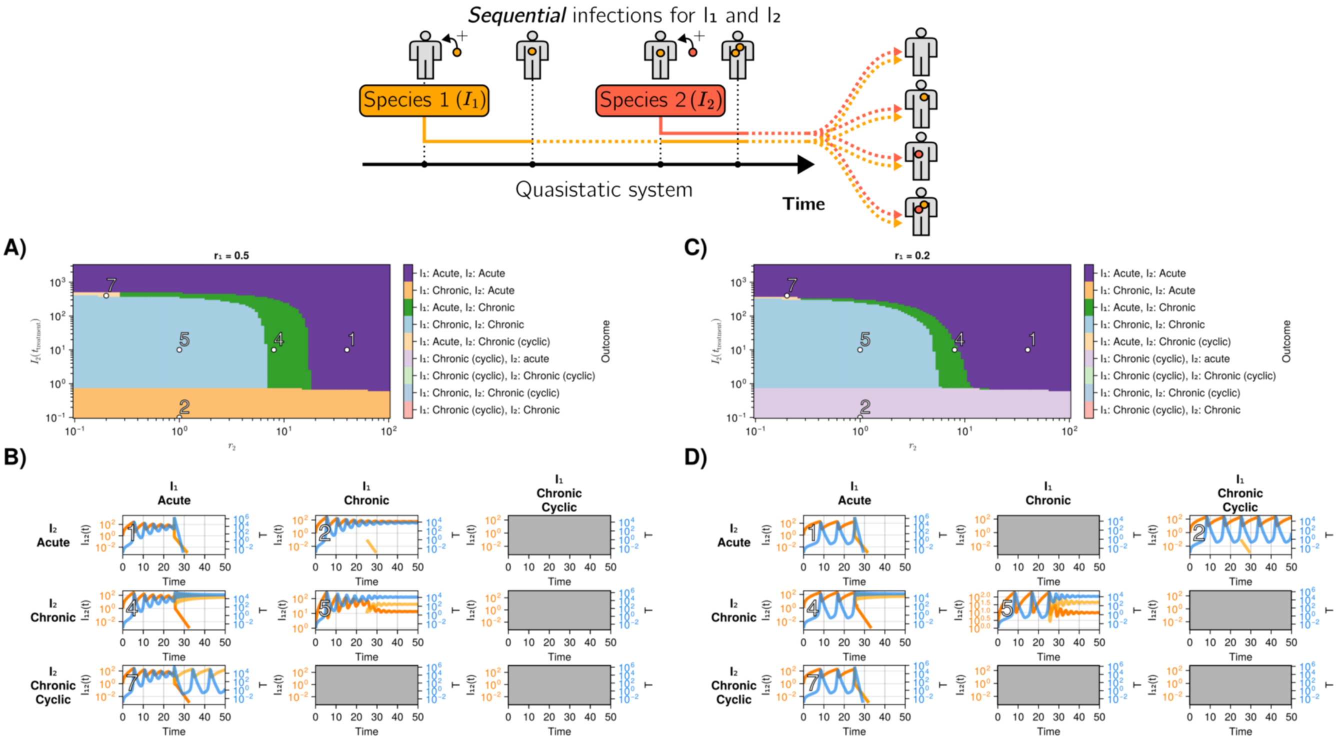


FIG. 13. A) Infection outcomes for the quasistatic system (equation 3) in the "treatment scenario", with initially a single species with $r_1 = 0.5$. It evolves towards a chronic state. The figure is to be compared with Fig. 6. At $t_{\text{treatment}} = 25$, a second species is added, with its own $r_2$, in amount $I\left(t_{\text{treatment}}\right)$. Examples of these processes are presented in sub-subplots B). B) Example time series associated with the labeled points in A). Some combinations are not attainable. C) Same as A), but the initial chronic state is in a cyclic state, with $r_1 = 0.2$. D) Example time series associated with the labeled points in C). Notice that some of the colors in the colorbar do not appear in the phase diagram.

point, is

$$\frac{\partial(\dot{T}_r, \dot{I}_r)}{\partial(T_r, I_r)}\Big|_{T_r^*, I_r^*} = \begin{pmatrix} \alpha_r I_r^* - 1 & \kappa_r + \alpha_r T_r^* \\ -I_r^* & r_r - T_r^* \end{pmatrix} . \tag{I4}$$

Solving for the eigenvalues $\lambda$ of this matrix, and writing everything in terms of $T_r^*$ by using the two forms of $I_r^*$ in eq. (I3), we find

$$\lambda_\pm = \frac{b}{2} \pm \frac{1}{2}\sqrt{b^2 - 4a} \quad \text{where} \tag{I5}$$
$$b = -\frac{\kappa_r}{\kappa_r + \alpha_r T_r^*} - (T_r^* - r_r) \quad \text{and} \quad a = \frac{\kappa_r + \alpha_r T_r^*}{T_r^* - r_r} .$$

We observe that $b < 0$ always, since it is a sum of two negative terms (recall that $T_r^* - r_r > 0$), so at least one eigenvalue has a negative real part. It turns out that both eigenvalues always have a negative real part, since $a > 0$. Indeed, there are two possibilities. 1) If $a$ is small, such that $b^2 > b^2 - 4a > 0$, then both $\lambda_\pm$ are real and negative, because the square root term's magnitude is not large enough to make $\lambda_+$ positive; in this case, the fixed point is a stable node. 2) If $a$ is large, then the discriminant is negative and $\lambda_\pm$ are a pair of conjugate complex eigenvalues with negative real part equal to $b < 0$; the point is then a stable spiral.

In practice, typical parameter values are $r_r \in [0.0, 4.0]$, $\kappa_r = 8 \times 10^{-8}$, $\alpha_r = 0.04$. This gives $a \in [0.04, 4.0]$ and $b \approx -0.04$, resulting in eigenvalues that are always imaginary with negative real part, i.e., a stable spiral (the damped oscillations mentioned in the main text).

### Appendix J: Timing of treatment

We have considered two cases for the addition of a second species. First, the case where both species are added simultaneously at the beginning of the system, and second, where one species reaches a chronic state and the second is subsequently added. As shown, the system can present some oscillations (transiently for the reduced system, and persistent for the full and quasistatic systems). It turns out that the timing of the addition of a second species, with respect to these oscillations, affects the outcome of an infection. For example, a second species added at the peak of the oscillations might not produce the same outcome as one added at the trough.

The effect the timing has can be understood using the notion of basins of attraction for a given amount of a second species, as shown in Fig. 5J. These basins are in reality cross-sections of a 3D "volume of attraction", in $I_1$, $I_2$, and $T$ space. A surface delimits this volume and contains the chronic fixed point. Hence, the addition of a second species will yield an acute reaction if its addition makes the trajectory cross this surface (hence exiting the volume of attraction).

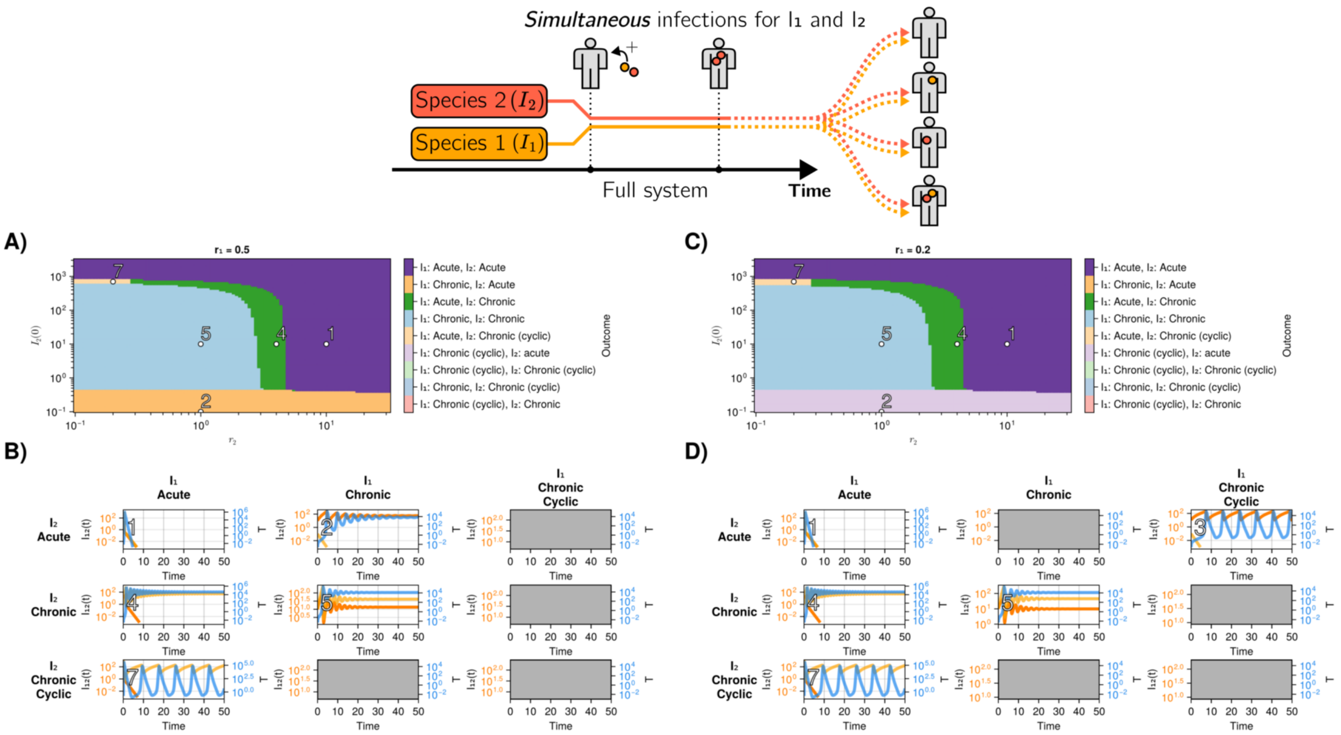


FIG. 14. A) Infection outcomes for the full system (equation 2) with two species initialized simultaneously in the system at $t = 0$ with $r_1 = 0.5$. The figure is to be compared with Fig. 5. Examples of these processes are presented in sub-subplots B). B) Example time series associated with the labeled points in A). Some combinations are not attainable. C) Same as A), but $I_1$ has $r_1 = 0.2$, which would produce a cyclic chronic infection if simulated alone. D) Example time series associated with the labeled points in C). Notice that some of the colors in the colorbar do not appear in the phase diagram.

Figure 18A shows a projection of this 3D space with an example trajectory. The black contour line delimits the cross-section of the volume of attraction for $I_2 = 330$. The addition of 330 of $I_2$ at the point B does not "dislodge" the trajectory from its volume of attraction, maintaining a chronic reaction (Fig. 18B). However adding it at point C will essentially be "piercing" across the basin and making the reaction acute (Fig. 18C). This explains how the timing of the addition of the second species can affect the system.

### Appendix K: Effect of $\gamma$

The $\gamma$ parameter controls the exhaustion or death rate of the T-Cells. It does not have a single variable analog in the reduced system since the variables $r_r$, $\kappa_r$ and $\alpha_r$ are all proportional to it. However, the system's dynamics do overall depend on it, as illustrated in Fig. 19.

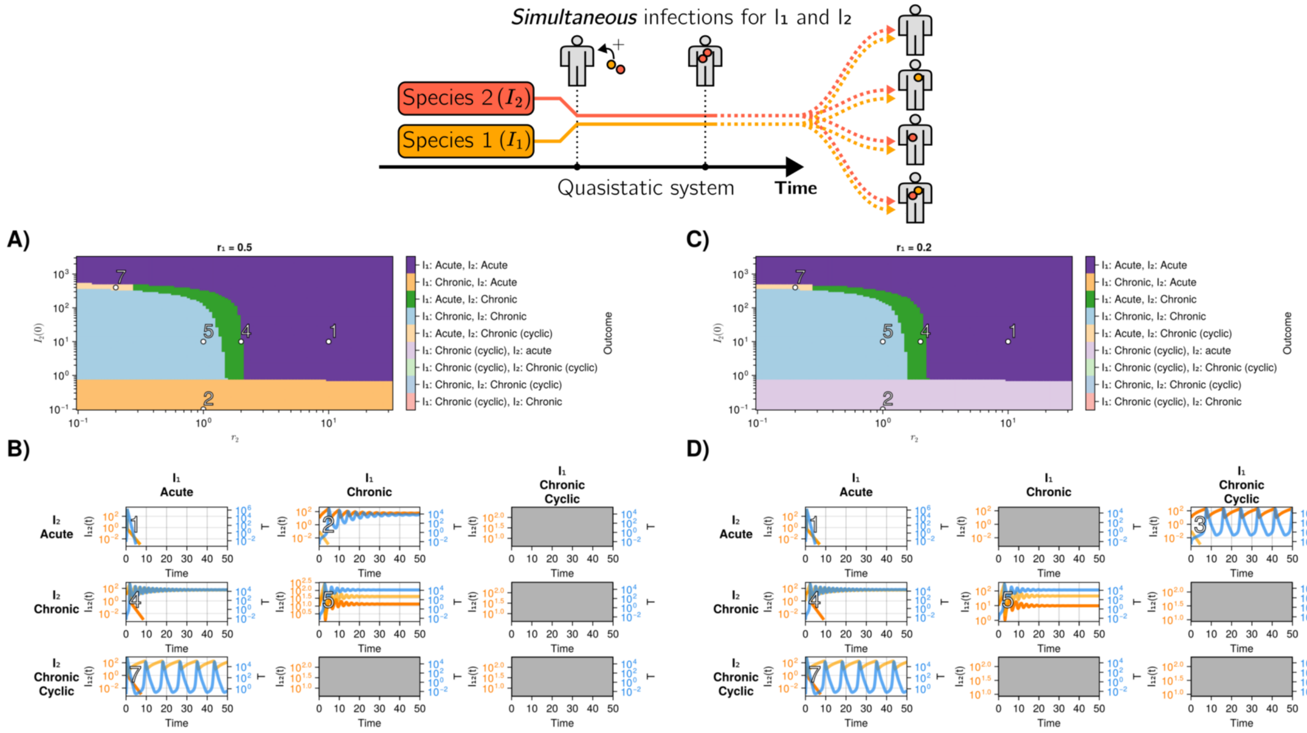


FIG. 15. A) Infection outcomes for the quasistatic system (equation 3) with two species initialized simultaneously in the system at $t = 0$ with $r_1 = 0.5$. The figure is to be compared with Fig. 5. Examples of these processes are presented in sub-subplots B). B) Example time series associated with the labeled points in A). Some combinations are not attainable. C) Same as A), but $I_1$ has $r_1 = 0.2$, which would produce a cyclic chronic infection if simulated alone. D) Example time series associated with the labeled points in C). Notice that some of the colors in the colorbar do not appear in the phase diagram.

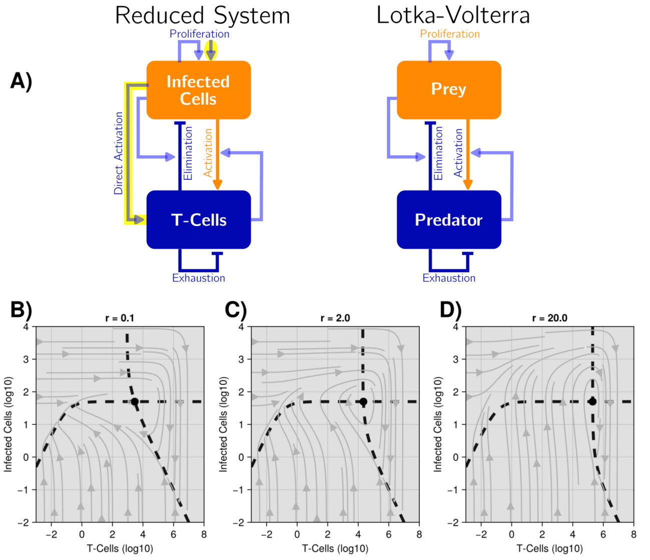


FIG. 16. A) Comparison between the reduced system of equation 1 and the Lokta-Volterra system. The differences are highlighted in yellow. B-D) Phase portrait of the reduced system without any threshold for infected cells. The system does not seem to undergo any bifurcations for positive values of $r$. Notice the basin of attraction spanning the entirety of the plot.

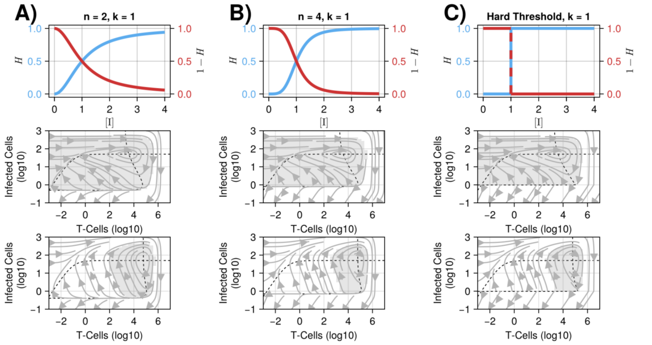


FIG. 17. A-C) Examples of the Hill functions used to interpolate between the two cases of equation H1 in equation H2. We show the resulting phase portraits, with their basins of attraction, for chronic and acute parameters for each case.

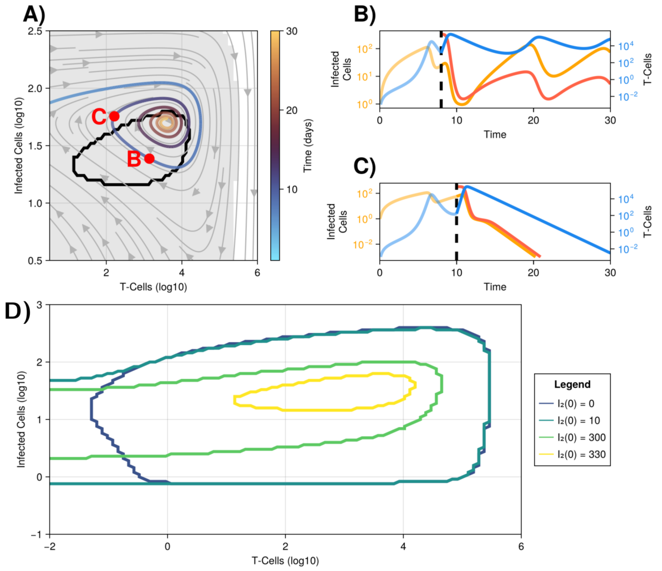


FIG. 18. The timing of the second infection affects the global outcome. A) Phase portrait of an infection with a single species. The black contour delimits the values of T-cells and infected cells for which a chronic infection will occur if 330 of a second species is added to the system. B) Chronic infection if the second species is added while the initial trajectory is inside the black contour. C) Acute infection if the second species is added while the trajectory is outside the black contour. D) Bassins of attraction for a challenge when different quantities of a second challenge are added.

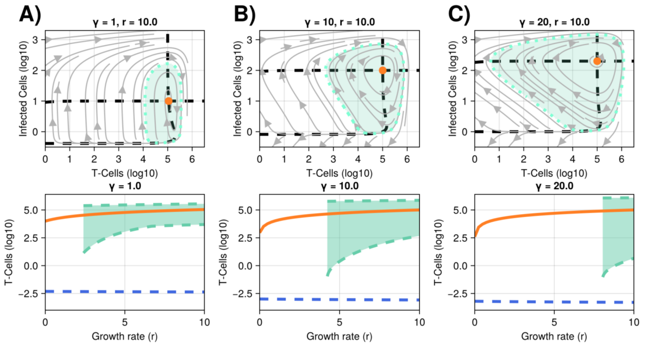


FIG. 19. The effect changing the $\gamma$ parameter has on the system. The unstable limit cycle shrinks for smaller $\gamma$. The boundary of the soft threshold gets lower. The location of the homoclinic bifurcation also shifts. The phase portraits are cross-sections of their corresponding bifurcation diagrams at $r = 10$. A) $\gamma = 1$. B) $\gamma = 10$. C) $\gamma = 20$.
